# Genome-culture coevolution promotes rapid divergence in the killer whale

**DOI:** 10.1101/040295

**Authors:** Andrew D. Foote, Nagarjun Vijay, María C. Ávila-Arcos, Robin W. Baird, John W. Durban, Matteo Fumagalli, Richard A. Gibbs, M. Bradley Hanson, Thorfinn S. Korneliussen, Michael D. Martin, Kelly. M. Robertson, Vitor C. Sousa, Filipe. G. Vieira, Tomáš Vinař, Paul Wade, Kim C. Worley, Laurent Excoffier, Phillip. A. Morin, M. Thomas. P. Gilbert, Jochen. B.W. Wolf

## Abstract

The interaction between ecology, culture and genome evolution remains poorly understood. Analysing population genomic data from killer whale ecotypes, which we estimate have globally radiated within less than 250,000 years, we show that genetic structuring including the segregation of potentially functional alleles is associated with socially inherited ecological niche. Reconstruction of ancestral demographic history revealed bottlenecks during founder events, likely promoting ecological divergence and genetic drift resulting in a wide range of genome-wide differentiation between pairs of allopatric and sympatric ecotypes. Functional enrichment analyses provided evidence for regional genomic divergence associated with habitat, dietary preferences and postzygotic reproductive isolation. Our findings are consistent with expansion of small founder groups into novel niches by an initial plastic behavioural response, perpetuated by social learning imposing an altered natural selection regime. The study constitutes an important step toward an understanding of the complex interaction between demographic history, culture, ecological adaptation and evolution at the genomic level.

---

The interplay between ecology, culture and evolution at the level of the genome remains poorly understood^1^. The ability to adapt to novel ecological conditions through behavioural plasticity is thought to be able to buffer natural selection pressures and promote rapid colonisation of novel niches^1^. However, by perpetuating exposure to a novel environment, stable cultural transmission of behaviour can also provide an opportunity for natural selection to act on adaptive genomic variation. Examples of genomic adaptation in humans during the period of recent ecological and cultural diversification and consequent demographic expansion are well illustrated^1-3^.

For example, the Inuit of Greenland descend from a small founder population that split from an East Asian source population and successfully colonised the extreme climatic conditions of the Arctic environment through culturally transmitted methods of hunting marine mammals and genetic adaptation to a cold climate and hypoglycemic lipid-rich diet^4,5^. However, our understanding of the complex interaction between ecology, culture, adaptation and reproductive isolation at a genome-wide level have long suffered from deficiency of genome-wide data, and conceptually, from the almost exclusive focus on these processes in humans and thus a lack of comparative data from other species^6^.

Killer whales *(Orcinus orca)* are the largest species in the dolphin family (Delphinidae) and together with humans, are one of the most cosmopolitan mammals, being found in all ocean basins and distributed from the Antarctic to the Arctic^7^. This top marine predator consumes a diverse range of prey species, including birds, fish, mammals and reptiles^7^. However, in several locations killer whales have evolved into specialized ecotypes with hunting strategies adapted to exploit narrow ecological niches^8-15^. For example in the North Pacific, two sympatric ecotypes co-exist in coastal waters: the mammal-eating (so-called *‘transient’*) ecotype and fish-eating (so-called *‘resident’*) ecotype^8-11^. This ecotypic variation is stable among multiple sub-populations of the *transient* and *resident* ecotypes across the North Pacific^8-11^ that diverged from common ancestral matrilines approximately 68 and 35 KYA respectively^16^. A highly stable matrilineal group structure and a long post-menopausal lifespan in killer whales is thought to facilitate the transfer of ecological and social knowledge from matriarchs to their kin^17^ and thereby perpetuate the stability of ecotypic variation in killer whales^18^. In the absence of a definitional consensus and for the purposes of investigating how cultural phenomena interact with genes, culture has been broadly defined as *‘information that is capable of affecting individuals’ behaviour, which they acquire from other individuals through teaching, imitation and other forms of social learning’*^1^. Several studies have argued that behavioural differences among killer whale ecotypes are examples of culture in this broader sense of the term^18,19^. However, this behavioural variation among ecotypes likely results from ecological, genetic and cultural variation and the interaction between them, rather than a single process explaining all behavioural variance^20^. Killer whales therefore offer a prime example of how behavioural innovation perpetuated by cultural transmission may have enabled access to novel ecological conditions with altered selection regimes, and thus provide an excellent study system for understanding the interaction between ecological and behavioural variation, and genome-level evolution.

## Results and Discussion

### Whole genome sequencing

We generated whole genome re-sequencing data of 48 individuals at low coverage and accessed high coverage sequencing data from two further individuals^21,22^ to investigate patterns of genomic variation among killer whale ecotypes. The samples represent five distinct ecotypes that, based on phylogenetic analysis of mitochondrial genomes^16^, include among the oldest and youngest divergences within the species (Fig. 1). The dataset incorporated 10 individuals each of the *transient* and *resident* ecotypes that occur in sympatry in the North Pacific; and from Antarctic waters, 7 individuals of a large mammal-eating form *(type B1)*, 11 individuals of a partially sympatric, smaller form which feeds on penguins *(type B2)*, and 10 individuals of the smallest form of killer whale, which feeds on fish *(type C)* (Fig. 1). A total of 2,577 million reads uniquely mapped to the 2.4-Gbp killer whale reference genome^21^ (Supplementary Fig. 1) for which a chromosomal assembly was generated for this study, so that approximately 50% of the autosomal regions of each individual were sequenced at ≥2× coverage (Supplementary Table 1). Subsequent data analyses used methods that account for uncertainty in the assignments of genotypes, enabling accurate inferences to be drawn from low-pass next-generation sequencing data^23,24^. Comparisons of estimated population genomic metrics such as genome-wide and per-site *F*_ST_ indicated that estimates from our low coverage data were highly consistent with published high coverage RAD-seq^25^ and SNP-typing^16^ data (Supplementary Fig. 2; Supplementary Table 2) and thus confirmed the robustness of our estimates.

**Figure 1.**
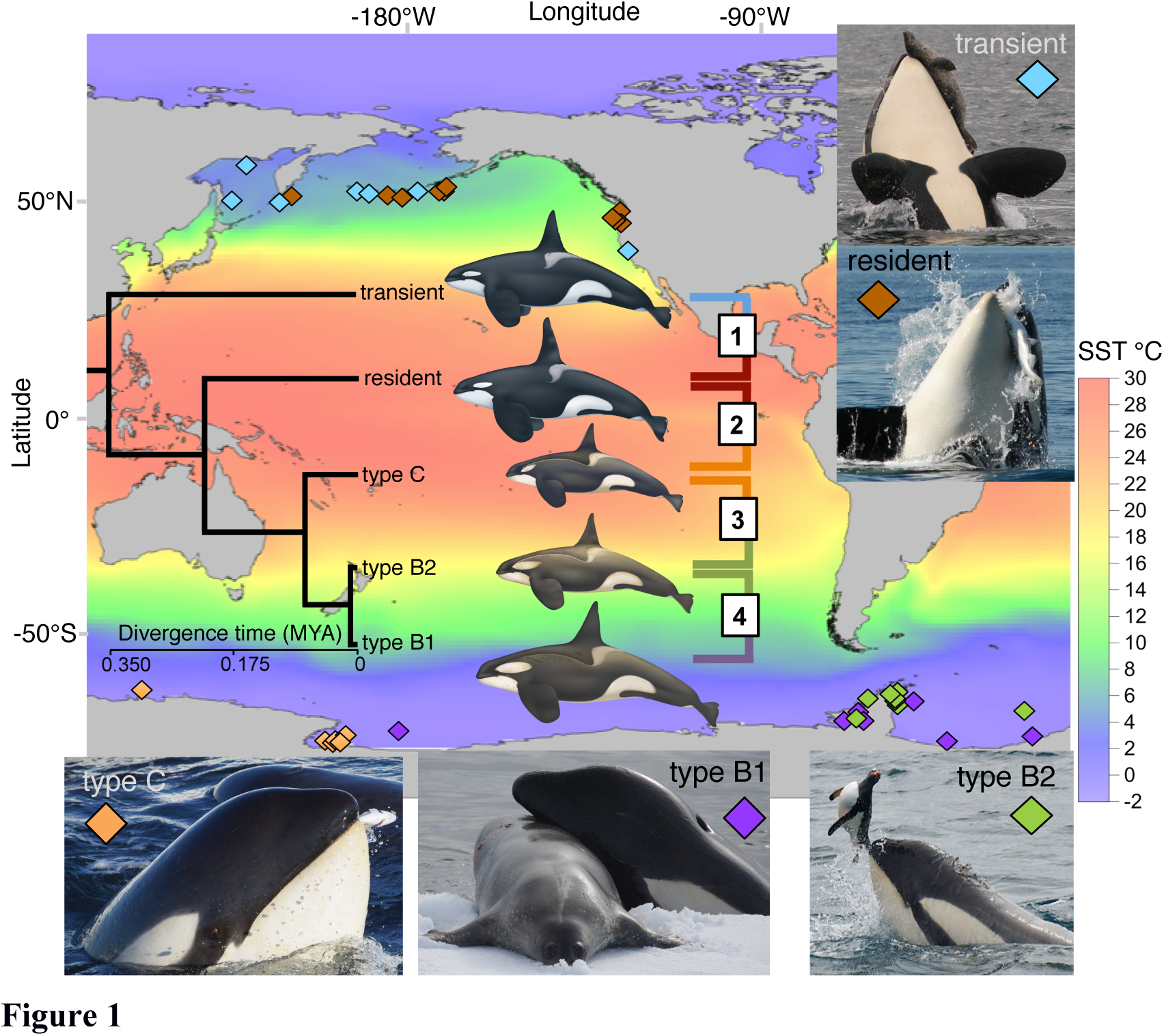
|Map of sampling locations of the five killer whale types included in this study. Sampling locations and inset photographs illustrating favoured prey species are colour coded by ecotype: *‘transient* (blue) and *type B1* (purple) are predominantly mammal-eating; *‘resident’* (brown) and *type C* (orange) are predominantly fish-eating; *type B2* (green) are known to feed upon penguins. The map is superimposed on a colour grid of sea surface temperature (SST). The Antarctic ecotypes primarily inhabit waters 8-16 °C colder than the North Pacific ecotypes. The relationship among these types and their estimated divergence times based on mitochondrial genomes are shown in the superimposed chronogram. Boxes 1-4 indicate pairwise comparisons spanning points along the ‘speciation continuum’ used to investigate the build up of genomic differentiation. [Photo credits: Dave Ellifrit, Center for Whale Research; Holly Fearnbach and Robert Pitman, SWFSC; SST measurements from NOAA Optimum Interpolation SST V2 long-term mean 1981-2010, www.esrl.noaa.gov/psd/repository courtesy of Paul Fiedler; killer whale illustrations courtesy of Uko Gorter.

### Time to most recent common ancestor (TMRCA)

We estimated a TMRCA of approximately 126–227 KYA from the accumulation of derived mutations at third codon positions for the most divergent killer whale lineages compared here (Supplementary Table 3) and based on the 95% highest posterior density interval (HPDI) of the mutation rate estimate^26^. This equates to approximately 4,900–8,800 generations and indicates a rapid diversification, over a timescale comparable to the diversification of modern humans^27^. We caution that for these age estimates we rely on mutation rate estimates derived from interspecific comparisons among odontocetes, and that therefore these estimates of TMRCA are at best approximate. Further, the demographic history and any gene flow between ecotypes will have an influence on the sharing of derived mutations and hence this estimate. However, we do expect that the estimate will be within the correct order of magnitude. Our estimated TMCRA overlaps with a recent RAD marker study^28^, which estimated a TMRCA of 189 KYA (only a point estimate was reported by these authors) scaling by a mutation rate 1.21 times higher than we have used here. The estimated TMRCA of a global dataset of killer whales based on non-recombining mitochondrial genomes has been estimated at 220-530 KYA^16^, older than our estimate based on nuclear genomes. Recombination of the nuclear genome among killer whale lineages has therefore continued after matrilineal lineages have diverged.

### Genetic differentiation and divergence

Despite this recent shared ancestry, substantial genome-wide differentiation and divergence had accrued between all pairs of ecotypes included in this study (Fig. 2, Supplementary Table 2). At *K*=5 populations, a maximum-likelihood–based clustering algorithm^29^ unambiguously assigned all individuals to populations corresponding to ecotype (Fig. 2c) indicating that all ecotypes have been assortatively mating long enough to allow allele frequencies to drift apart. Pairwise genetic distances between individuals visualized as a tree indicate that segregating alleles are largely shared within an ecotype (Supplementary Fig. 3). Similarly, pairwise relatedness due to identity-by-descent (IBD), *i.e.* genetic identity due to a recent common ancestor was high within each ecotype. While the three Antarctic ecotypes still showed signs of recent relatedness, no shared recent IBD ancestry was detected between Antarctic and Pacific types or between the sympatric *resident* and *transient* ecotypes (Supplementary Fig. 4). The greatest differentiation (Supplementary Table 2) as visualised in the maximum likelihood graph (Fig. 2a) and PCA plot (Fig. 2b) was between the allopatric Pacific and the Antarctic ecotypes, whilst differentiation among Antarctic ecotypes was much lower than between the two Pacific ecotypes. Thus, our sampled populations allowed us to investigate the accrual of genomic differentiation along points of a continuum, acting as a proxy of sampling at different stages of the speciation process^30^. The accrual of genome-wide differentiation *(F*_ST_ = 0.09) between even the most recently diverged and partially sympatric ecotypes (Antarctic types *B1* and *B2*) indicates that reproductive isolation quickly becomes established after the formation of new ecotypes. Thus whole-genome resolution confirms that even in sympatry, contemporary gene flow occurs almost exclusively among individuals of the same ecotype, allowing genomic differentiation to build up between ecotypes so that within an ocean basin, ecological variation better predicted genetic structuring than geography.

**Figure 2.**
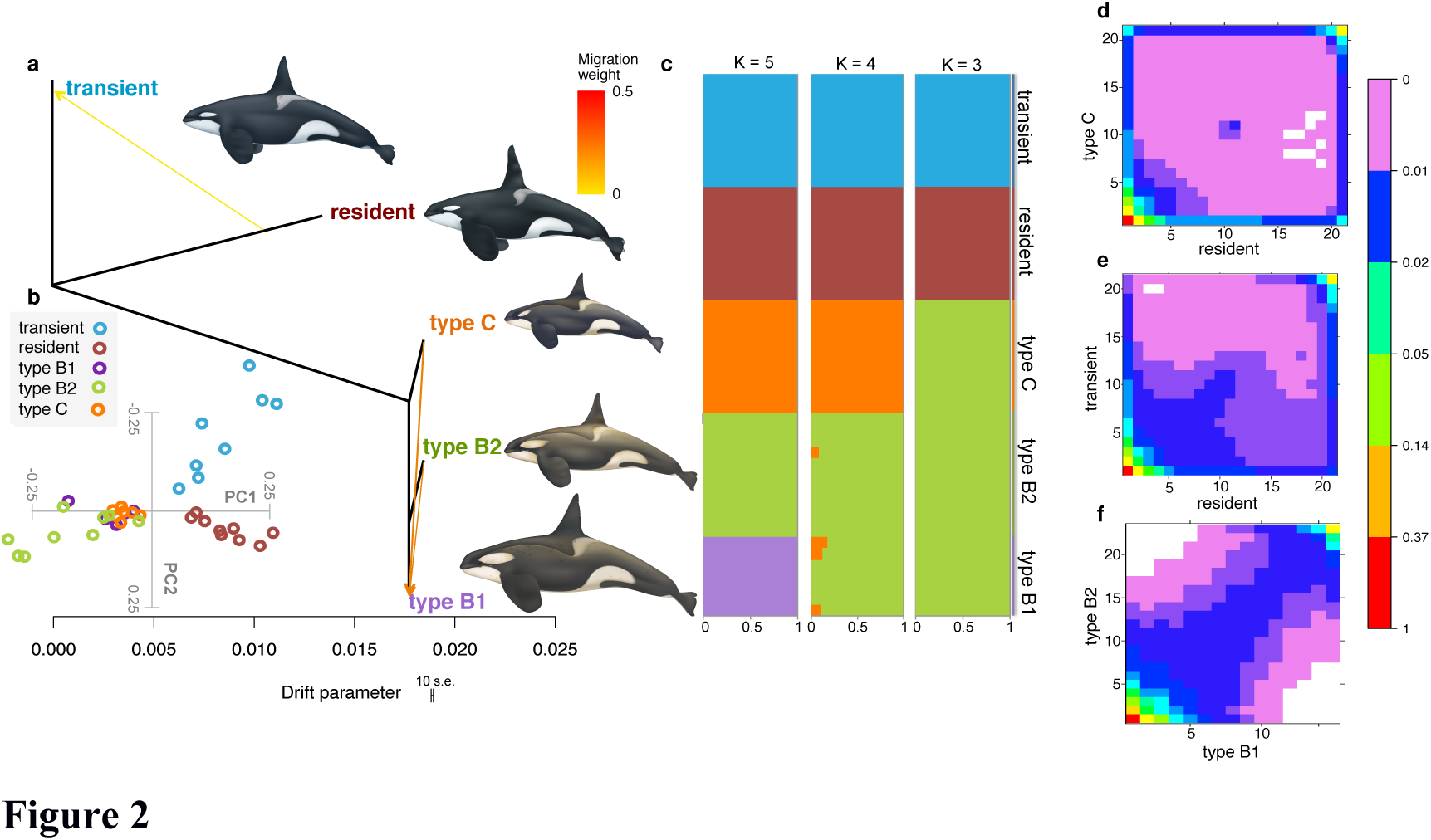
|Evolutionary relationships among killer whale ecotypes. **(a)** TreeMix maximum likelihood graph from whole-genome sequencing data, rooted with the *‘transient’* ecotype. Horizontal branch lengths are proportional to the amount of genetic drift that has occurred along that branch. The scale bar shows ten times the average standard error of the entries in the sample covariance matrix. Migration edges inferred using TreeMix and supported by the *f*3 statistic test are depicted as arrows coloured by migration weight, **(b)** Principal Component Analysis (PCA), both the first and second principal components were statistically significant (p-value < 0.001). **(c)** Ancestry proportions for each of the 48 individuals conditional on the number of genetic clusters *(K*=3-5). **(d-f)** Joint site-frequency spectra for pairwise comparisons illustrating the speciation continuum. Each entry in the matrix (*x,y*) corresponds to the probability of observing a single-nucleotide polymorphism (SNP) with frequency of derived allele *x* in population 1 and *y* in population 2. The colours represent the probability for each cell of the site frequency spectrum (SFS), white cells correspond to a probability of zero. The analysis illustrates that the amount of genetic differentiation is greater between **(d)** Pacific and Antarctic populations, and **(e)** the pair of Pacific types than **(f)** among the evolutionarily younger Antarctic types, which have highly correlated site frequency spectra.

### Ancient admixture

To better understand and visualise the complexity of the ancestry of killer whale ecotypes we reconstructed the genetic relationships among ecotypes in the form of a maximum likelihood graph (Fig. 2a) representing the degree of genetic drift and modelling both population splits and gene flow using the unified statistical framework implemented in TreeMix^31^. The inferred migration edges were supported by the three-population (*f*3) and D-statistic (ABBA-BABA) tests, which can provide clear evidence of admixture, even if the gene flow events occurred hundreds of generations ago^32-34^. These population genomic methods test for asymmetry in the covariance of allele frequencies that indicate the relationships among populations are not fully described by a simple bifurcating tree model.

The three approaches were consistent in inferring migration from source populations that share ancestry with the North Pacific *resident* and North Atlantic ecotype into the *transient* ecotype (Fig. 2a, Supplementary Figs. 5 & 6, Supplementary Tables 4-6). The genomes of *transients* are therefore partly derived from at least one population related to the Atlantic and *resident* ecotype, (but not necessarily these populations, *i.e.* the source could be an un-sampled ‘ghost’ population). The asymmetrical 2D-site frequency spectrum (SFS) also implies directional gene flow from a population ancestrally related to the *residents* into the *transient* ecotype^35^ (Fig. 2e). Given the extent of the inferred demographic bottlenecks during founder events (see section below), and the expected consequential shift in the SFS^36,37^, it seems likely that this admixture would have occurred during or after the founder bottleneck in the *transient* ecotype, otherwise it would be expected to be less correlated with the *resident* SFS. The sequencing of more populations is expected to shed further light on this episode of ancient admixture. Previous studies have inferred on-going gene-flow between the *resident* and *transient* ecotypes^38,39^, however, the outcome of the suite of analyses applied here strongly imply reproductive isolation between present-day populations (Fig. 2).

TreeMix, the three-population and D-statistic tests inferred that type *B1* is admixed and derives from at least two populations related to both types *B2* and *C* (Fig. 2a, Supplementary Tables 4-6). Conducting D-statistic tests on proposed tree-like histories comprising combinations of 16 genome sequences that included the North Atlantic sequence, we found that Antarctic *types B1* and *B2* shared an excess of alleles with the three Northern Hemisphere ecotypes (Supplementary Figs. 7 & 8). This shared ancestry component between Northern and Southern hemisphere ecotypes was not detected in *type C*, suggesting a relatively recent admixture event after *type C* split from the shared ancestor of *types B1* and *B2*. Other signals of ancient admixture among populations were also detected and are reported in the tables in the Supplementary Material.

### Demographic history

As our results suggest that killer whale ecotypes have diversified rapidly from a recent ancestor and given the importance of the relationship between effective population size (*N*_e_) and the rate of evolution^40^, we conducted analyses to reconstruct their demographic history. Applying the pairwise sequential Markovian coalescent (PSMC) approach^27^ to two high coverage (≥20×) autosomal assemblies, a North Atlantic female and a North Pacific *resident* male, refining the methodological approach of a previous analysis of these genomes (Supplementary Figs. 9 & 10), we recovered a similar demographic trajectory to that previously reported^22^ (Fig. 3a). The inference of the timing of these demographic declines is dependent upon the assumed mutation rate (*μ*), but across a range of sensible estimates of *μ*, the declines broadly fall within the Late Pleistocene^22^. This was previously interpreted as evidence for demographically independent population declines in each ocean, driven by environmental change during the Weichselian glacial period^22^. However, this inference assumes that each PSMC plot tracks the demographic history of a single unstructured panmictic population^41^. The y-axis of the PSMC plot is an estimation of *N*_e_ derived from the rate of coalescence between the two chromosomes of a diploid genome. However, in the presence of population structuring, regions of the two chromosomes will coalesce less frequently as their ancestry may derive from different demes or sub-populations; the rate of coalescence can thus be similar to a single population with large *N*_e_. PSMC estimates of *N*_e_ during population splits can therefore be greater than the sum of the effective sizes of the sub-populations, dependent on the number of sub-populations and the degree of connectivity *(i.e.* cross-coalescence) between them^27,41^ (see Figure S5 of ref 27). In fact, the results presented here and previously published^16,25,28^ indicate that throughout the Weichselian glacial period there were multiple population splits, both between and within ecotypes, including the splitting of the two lineages included in the PSMC analyses just at the point of the change in inferred *N*_*e*_ (Fig. 3a; Supplementary Fig 11). Therefore, the PSMC plots will be strongly influenced by these changes in structure, and the changes in inferred *N*_e_ are not necessarily associated with population declines (see Supplementary Fig. 12).

**Figure 3.**
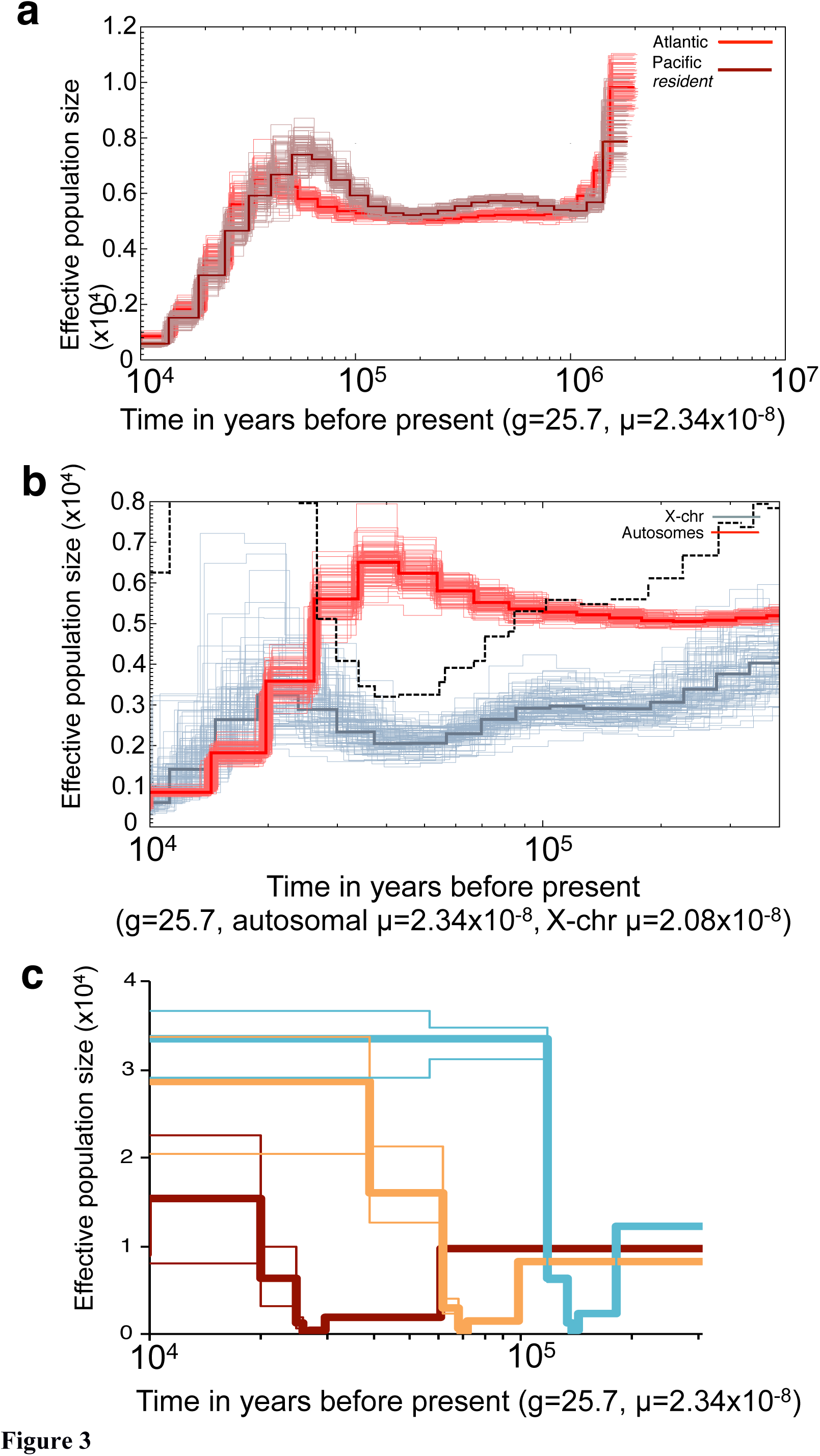
|Reconstructing the demographic history of killer whale ecotypes. **(a)** Pairwise sequential Markovian coalescent (PSMC) estimates of changes in effective population size (*N*_e_) over time inferred from the autosomes of a North Atlantic killer whale (red) and from the autosomes of a North Pacific *resident* killer whale (brown). Thick lines represent the median, thin light lines of the same colour correspond to 100 rounds of bootstrapping, **(b)** PSMC estimates of changes in *N*_e_ over time inferred from the autosomes (*N*_eA_, red) and the X-chromosome (*N*_ex_, grey) of the high coverage genome sequence of a North Atlantic female killer whale. Thick lines represent the median, thin light lines of the same colour correspond to 100 rounds of bootstrapping. The dashed black line indicates the ratio of *N*_eX_/*N*_eA_ **(c)** Changes in effective population size (*N*_e_) over time in the *transients* (blue), *residents* (brown) and type C (orange) inferred using the site frequency spectrum (SFS) of each ecotype. Thick lines represent the median, thin light lines the 2.5 and 97.5 percentiles of the SFS analysis.

To investigate the influence of connectivity on our PSMC plots further, we inferred ancestral *N*_e_ of the diploid X-chromosome (*N*_eX_) of the Atlantic female and directly compared to the autosomal fraction (*N*_eA_) of the genome presented in figure 3a. From around 300,000 to 130,000 years BP during the Saalian glacial period, the inferred *N*_eX_ ranged from 0.58-0.79 of the *N*_eA_ (Fig. 3b), which lies within expectation for demographically stable mammalian species^42^. During the first part of the Weichselian glacial period, *N*_eX_ markedly declined, reaching a minimum at approximately 30,000-50,000 years BP. The timing of this bottleneck in *N*_eX_ overlaps with the stem age of the mitochondrial clade for this Atlantic population^16^, *i.e.* consistent with almost all mitochondrial diversity being lost in this lineage during this period. Conversely, this is concurrent with the peak estimate of autosomal *N*_eA_ inferred by PSMC, and ratio of *N*_eX_/*N*_eA_ falls to approximately 0.3 at this *N*_eX_ minimum and then recovers within 1,000 generations to >0.75 (Fig. 3b). Simulated demographic bottlenecks of several hundred-fold reduction result in a disproportionate loss of *N*_eX_ attributed to the difference in inheritance mode of each marker, and the ratio of *N*_eX_/*N*_eA_ can reach less than 0.3^42^. Following the bottleneck, *N*_eX_ recovers more rapidly than *N*_eA_ and the ratio of *N*_eX_/*N*_eA_ can exceed 0.75 during this recovery phase^42^. The concordance of the timing of the bottleneck in the X-chromosome and the stem age of the mitochondrial genome suggest an underlying demographic process, rather than strong selection on the X-chromosome or some other factor driving mutation rate^42^. A sex-biased process such as primarily male-mediated gene flow between demes could further influence the ratio of *N*_eX_/*N*_eA_^43^.

To estimate the demographic histories from our population genomic data we produced ‘stairway’ plots using composite likelihood estimations of theta (*θ*) for different SNP frequency spectra associated with different epochs, which are then scaled by the mutation rate to estimate *N*_e_ for each epoch^44^. Using this method we reconstructed a demographic history from our population genomic data for the *resident* ecotype that was comparable to the PSMC plot from a single *resident* high coverage genome, both methods identifying a decline in *N*_e_ starting at approximately 60 KYA (Fig. 3a,c). The stairway plots for the *transient* ecotype and *type C* showed the same pattern as the resident, of a decline to a bottlenecked population with an *N*_e_ of <1,000, and a subsequent expansion (Fig. 3c). The bottlenecks did not occur simultaneously, as might be expected in response to a global environmental stressor during a glacial cycle, but instead were sequential (Fig. 3c). In each case, the timing of the demographic bottleneck overlapped with the previously estimated timing of the stem age of the mitochondrial genome clades containing each ecotype^16^. Thus, within the *transient, resident* and combined Antarctic ecotypes, both mitochondrial and nuclear lineages coalesce back to these bottleneck events, consistent with genetic isolation of small matrifocal founder groups from an ancestral source population, followed by the subsequent expansion and sub-structuring of a newly established ecotype.

Overall, the population genetic analyses of whole genome sequences above shed light on the ancestry of killer whale ecotypes in unprecedented detail, highlighting a complex tapestry of periods of isolation interspersed with episodic admixture events and strong demographic bottlenecks in the founder populations of the resident, transient and ancestral Antarctic ecotypes.

### Genome-wide landscape of genetic diversity and differentiation

Demographic bottlenecks during population splits and founding events, followed by subsequent demographic and geographic expansion can produce rapid shifts in allele frequencies between populations^3,45^. The high levels of genome-wide differentiation (*F*_S_T) between killer whale ecotypes across all genomic regions (Fig. 4a,b) are consistent with strong genetic drift following demographic expansion from small founding groups. Considering the low efficiency of selection in populations as small as the estimates presented here^46^, in which founder populations have an estimated *N*e ranging from a few tens to hundreds, a genome-wide contribution of ecologically-mediated divergent selection is neither necessary, nor particularly likely to explain the observed shifts in allele frequencies in such a large number of loci. Consistent with this prediction, we find that differentiation is highest along the branches inferred by TreeMix to have experienced the most substantial genetic drift (Fig. 2a), *i.e.* the branch to the ancestor of the Antarctic types and the branch to the resident ecotype (Supplementary Fig. 13). We therefore expect that only those beneficial alleles that have a strong favourable effect (i.e. strength of selection *(s*) > 1/2*N*e) would have an increased fixation probability due to selection within these founder populations.

**Figure 4.**
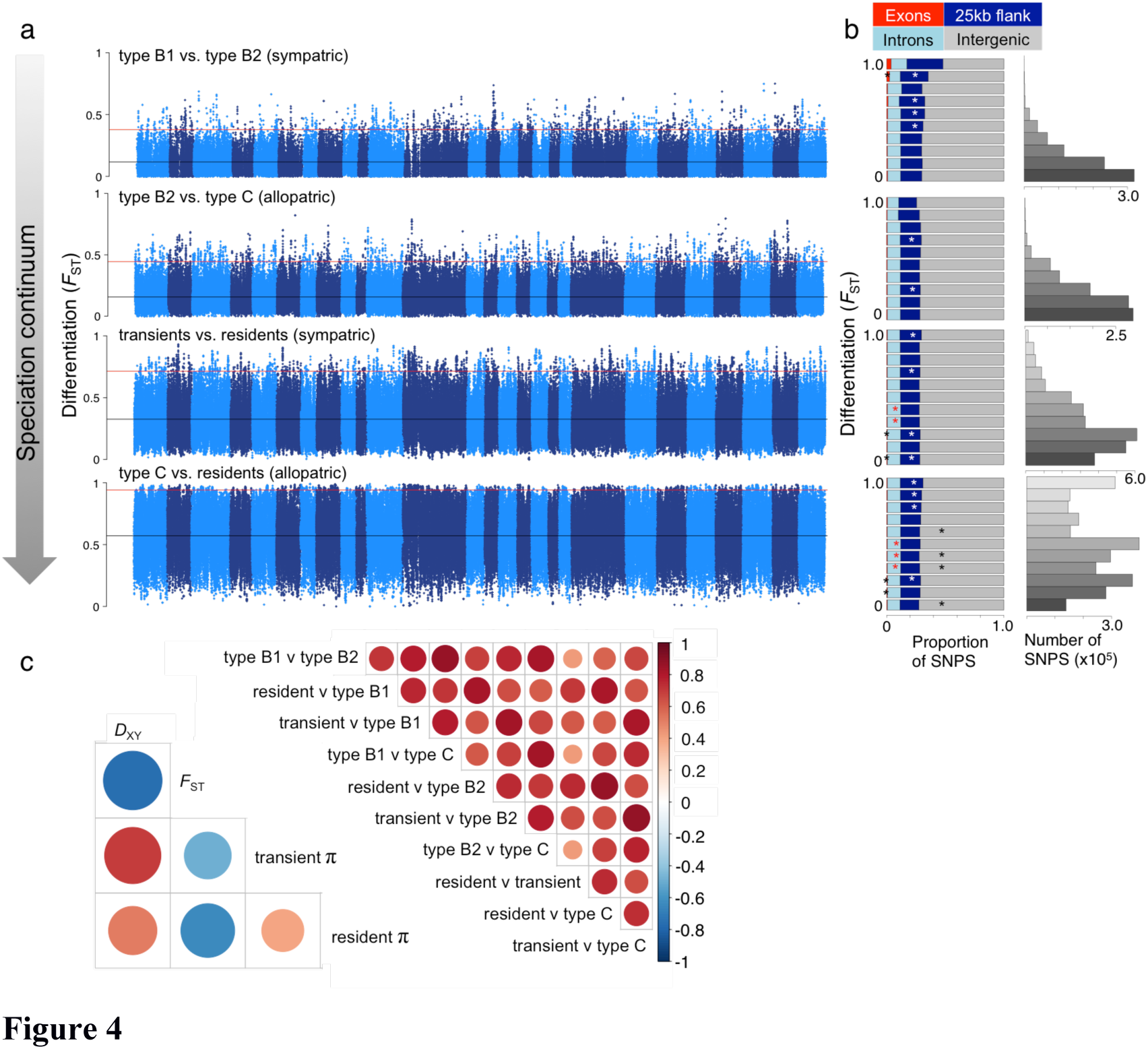
|Genome-wide distribution of differentiation. **(a)** Pairwise genetic differentiation (FST) in 50-kb sliding windows across the genome between killer whale ecotypes. Pairwise comparisons are arranged from the youngest divergence between sympatric ecotypes to older divergences of now allopatric ecotypes. Alternating shading denotes the different chromosomes; the horizontal black lines mark the mean *F*_ST_ and the horizontal red lines mark the 99th-percentile *F*_ST_ **(b)** Stacked bar plots show the proportion of SNPs in different genomic regions: exons, introns, 25-kb flanking and intergenic regions, in different *F*_ST_ bins, corresponding to the y-axis of the Manhattan plots for each pairwise comparisons. Asterisks signify an over-representation of SNPs in a given region at each *F*_ST_ bin. Bar plots indicate the total number of SNPs in each *F*_ST_ bin for each pairwise comparison, **(c)** Correlations of 50-kb window-based estimates of differentiation (*F*_ST_) between all possible pairwise comparisons between ecotypes (above diagonal); correlations between 50-kb window-based estimates of genome-wide divergence (*D*_XY_), differentiation (*F*_ST_) and nucleotide diversity *(π)* for the comparison between the resident and transient ecotypes (below the diagonal), other comparisons are shown in Supplementary Fig. 13. Red indicates a positive relationship, blue a negative one; colour intensity and circle size are proportional to Spearman’s correlation coefficient. Regions of high differentiation, but low diversity and divergence that are shared across pairwise comparisons are likely to have been regions of selection on the ancestral form, which would remove linked neutral diversity and result in increased lineage sorting of allele frequencies in these genomic regions in the derived forms.

Much of the heterogeneity in the differentiation landscape was shared among pairwise comparisons (Fig. 4c). Since diversity (*π*) was not associated with mutation rate μ, as inferred from neutral substitution rate *d*S, (*r* = −0.1 to −0.24), the observed co-variation in differentiation (*F*_ST_), diversity (*π*) and absolute divergence (*D*_xy_) between population pairs (Fig. 4c, Supplementary Fig. 14) is best explained by the shared local reduction of diversity by linked selection in the ancestral population^47,48^. This leads us to conclude that overall the landscape of genome-wide differentiation is a result of global genetic drift, regionally elevated by ancestral linked selection (e.g. background selection or repeated selective sweeps shared among populations) independent of the evolutionary dynamics of the recently derived present-day ecotypes.

### Genomic signatures of climate and diet adaptation

Yet, against this background of shared differentiation, there was evidence for genic divergence of putative functional relevance. The first targets of selection following ecotype diversification and the exploitation of new ecological niches are expected to be those that facilitate ecological specialization^47^. Once a certain level of reproductive isolation is reached, differences can accumulate in other (fast evolving) genes involved in reproductive isolation effectively reducing hybrid mating^49^. Following this rationale both individual gene associations and gene ontology enrichment analyses yielded several biological processes and candidate genes with putative functional roles in ecological specialization, local adaptation and reproductive isolation (Supplementary Tables 7-9). For example, in comparisons between ecotypes inhabiting the extreme cold of the Antarctic pack ice with ecotypes from the more temperate North Pacific (Fig. 1) we found the most significant enrichment in genes involved in adipose tissue development (GO:0060612, Fisher’s exact test: *P* = 0.0015). Genes associated with adipose tissue development have previously been found to be evolving under positive selection in the polar bear^50^, suggesting a role for this process in rapid adaptation to a cold climate and lipid-rich diet.

Using the population branch statistic (PBS), which has strong power to detect recent natural selection^5,51^ and allowed us to investigate allele changes along specific branches, we identified another candidate example, where cold-adaptation may play a role. The *FAM83H* gene showed a signature of selection (top 99.9% PBS values) and was found to contain four fixed non-synonymous substitutions derived in the Antarctic lineages, based on the inferred ancestral state, which resulted in physicochemical changes including a hydrophobic side chain being replaced by a positively charged side chain. The keratin-associated protein encoded by the *FAM83H* gene is thought to be important for skin development and regulation^52,53^, through regulation of the filamentous state of keratin within cytoskeletal networks in epithelial cells, determining processes such as cell migration and polarization^54,55^. Skin regeneration is thought to be constrained in killer whales whilst inhabiting the cold waters around Antarctica due to the high cost of heat loss, and is thought to underlie rapid round-trip movements to warmer subtropical waters by Antarctic ecotypes^56^. The balance between skin regeneration and thermal regulation in Antarctic waters could be a major selective force requiring both behavioural^56^ and genomic adaptation (Supplementary Fig. 15).

Genes encoding proteins associated with dietary variation also showed a signature of selection (top 99.9% PBS values). For example, the *carboxylesterase 2 (CES2)* gene encodes the major intestinal enzyme and has a role in fatty acyl and cholesterol ester metabolism in humans and other mammals^57^. Two exons of the *CES2* gene had among the top 99.9 percentile PBS values due to changes in allele frequencies (including 2 fixed non-synonymous amino acid changes) along the branch to the Antarctic types (Supplementary Fig. 16). Similarly, genes in the top 99.9 percentile PBS values were enriched for carboxylic ester hydrolase activity (GO:0052689, Fisher’s exact test: *P* < 0.0001) in the fish-eating *resident* ecotype. Biological processes enriched in the resident killer whale included digestive tract morphogenesis (GO:0048546, Fisher’s exact test: *P* = 0.0022), and gastrulation with mouth forming second (GO:0001702, Fisher’s exact test: *P* = 0.0024): associated with the formation of the three primary germ layers of the digestive system during embryonic development. These results overlapped with enriched GO-terms identified by a previously published RAD-seq study, despite the relatively sparse sampling of 3,281 SNPs in that study^25^. Enrichment of these GO-terms was largely driven by differentiation in a single exon in the *GATA4* gene, which included a fixed non-synonymous substitution, sequenced in both studies. Nonetheless this overlap highlights that reduced representation libraries can provide useful genome-wide inference of candidate targets of selection.

Signatures of selection along branches leading to the two predominantly mammal-eating ecotypes included in this study, the North Pacific *transient* and Antarctic *type B1*, were found in genes that play a key role in the methionine cycle (Fig. 5). Methionine is an essential amino acid that has to be obtained through dietary intake, and is converted through trans-sulfurcation to cysteine via intermediate steps of catalysis to homocysteine^58^. Any excess homocysteine is re-methylated to methionine^58^. Diets with different protein content, such as between killer whale ecotypes, will differ in their content of methionine, and the enzymatic cofactors involved in the metabolism of methionine and homocysteine, which include folate, vitamins B6 and B12^59^ (hence why vegetarians often take vitamin B12 supplements). Whilst different genes and different biological processes showed a signature of selection in each of these two mammal-eating ecotypes (Fig. 5), in both cases the candidate genes and processes were associated with the regulation of methionine metabolism, which results in the generation of cysteine. Successful hunting of mammal prey by killer whales would provide a sudden and rich source of dietary methionine. This fluctuating intake of protein may place more of a selective pressure on the regulation of the metabolism of methionine than does the consumption of fish by piscivorous ecotypes.

**Figure 5.**
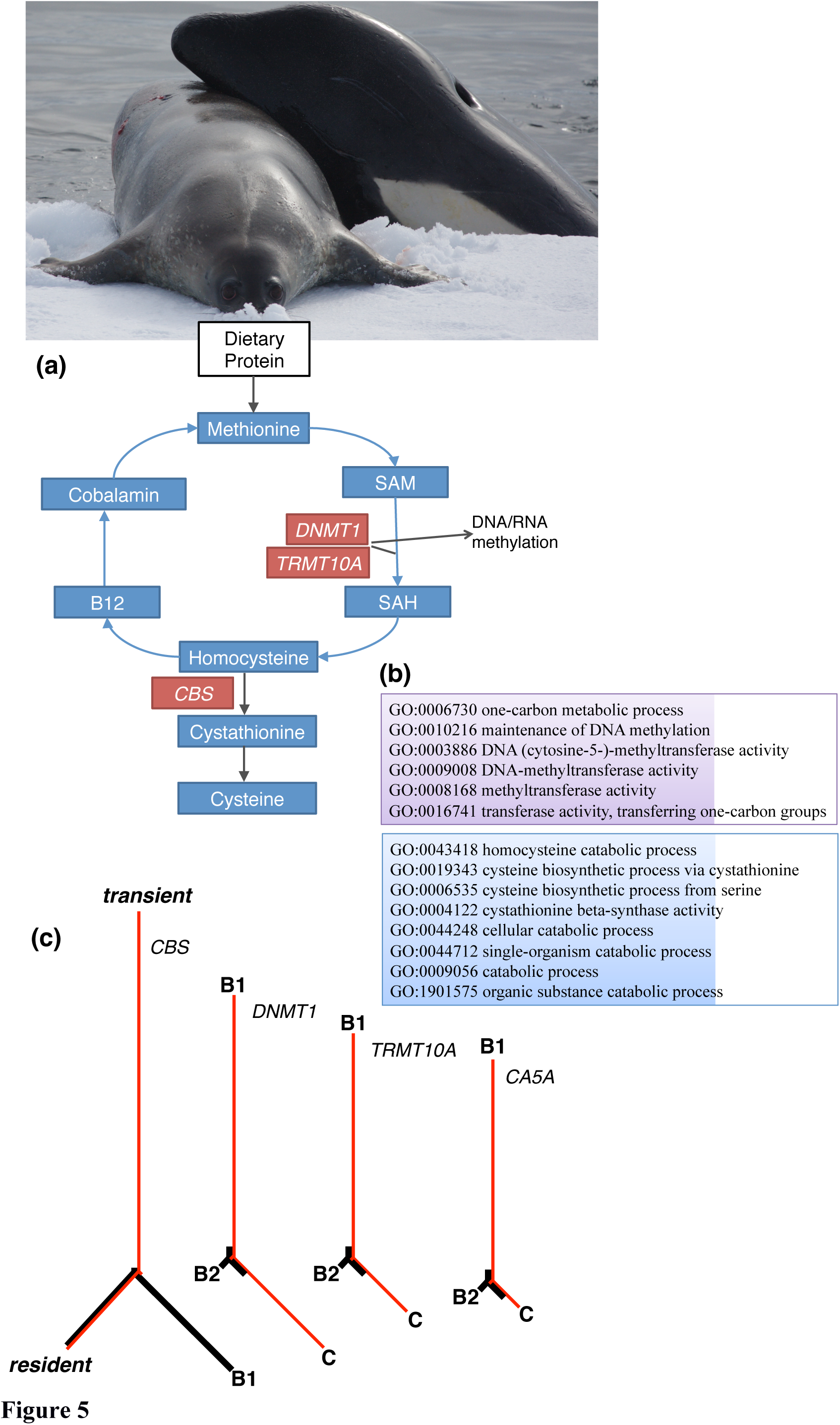
|Signatures of selection in mammal-eating ecotypes in genes that play a key role in the methionine cycle. **(a)** Methionine is an essential amino acid that is obtained through dietary protein. The methionine cycle feeds into the folate cycle and both are part of the complex and interacting network of pathways that encompass one-carbon metabolism^58^. Methionine adenyltransferase (MAT) then generates S-adenosylmethionine (SAM), which is then demethylated by the methyltransferases *DNMTl*^99^ and *TRMT10A*^100^ to methylate DNA and RNA respectively, and to form S-adenosylhomocysteine (SAH)^99^. SAH is converted to homocysteine, which is then catalysed by cystathionine P-synthase, an enzyme encoded by the *CBS* gene to cystathionine, which is then converted to cysteine^99^. Any excess homocysteine is remethylated to methionine. Adapted and simplified from KEGG pathway map 00270. Genes that play a role in this pathway and with a signature of selection (top 99.9% PBS values) in either of the predominantly mammal-eating ecotypes (the *transient* or *type B1)* are in red boxes. The gene encoding the carbonic anhydrase *CA5A*, which plays a role in the larger one-carbon pathway also had a signature of selection (top 99.99 PBS value) in *type B1.* **(b)** Boxes indicate GO terms associated with the biological processes within the methionine cycle that were enriched in genes with the top 99.9 *(transients;* blue) and 99.99 *(type B1;* purple) percentile PBS values, **(c)** Evolutionary trees underlying the signal of selection. Population-specific allele frequency changes are indicated by the FsT-based (PBS values) branch lengths in these genes (red), which are overlaid on the genome-wide average branch lengths (black). The high (top 99.9-99.99%) PBS values indicate substantial changes in allele frequencies along the branches to the mammal-eating ecotypes. Differentiation in genes DNMTl, TRMT10A and CA5A was greatest between the mammal-eating type B1 and the fish-eating type C among the Antarctic types. Photo credit of ‘dietary protein‘: Robert Pitman, SWFSC.

### Rapid evolution of reproductive proteins

High (top 99.9 percentile) PBS values, largely driven by fixed non-synonymous amino acid substitutions, were further estimated for several genes that encode proteins associated with reproductive function, including testes development, regulation of spermatogenesis, spermatocyte development and survival, and initiating the acrosome reaction of the sperm (e.g., *PKDREJ, RXFP2, C9orf24, SPEF1, TSSK4, DHH, MMEL1).* Reproductive proteins such as *PKDREJ*, in which we found two fixed non-synonymous substitutions derived in the ‘resident’ ecotype, are known to diverge rapidly across taxa, and due to their functional role in fertilization, are emerging as candidates for the postzygotic component of the speciation process^49,60^.

## Conclusions

Overall, our results indicate that the processes underlying genomic divergence among killer whale ecotypes reflect those described in humans in several respects. Behavioural adaptation has facilitated the colonization of novel habitats and ecological niches. Founder effects and rapid formation of reproductive isolation, followed by population expansion have promoted genome-wide shifts in the frequency of alternative alleles in different ecotypes due to genetic drift. Demographic changes during founder events and subsequent expansions can also influence cultural diversity^61,62^, and may have had a role in reducing within-ecotype cultural diversity and promoting cultural differentiation between ecotypes. As with studies on modern humans, it is difficult to demonstrate a causal association between cultural differences and selection on specific genes^1^, but our findings of divergence in genes with putative functional association with diet, climate and reproductive isolation broadly imply an interaction between genetically and culturally heritable evolutionary changes in killer whale ecotypes. Given these findings, the almost exclusive focus on humans by studies of the interaction of culture and genes^6^ should be expanded, and exploration of culture-genome coevolution models in suitable non-human animal systems encouraged.

## METHODS

### Sample collection

Skin biopsies from free-ranging killer whales were collected using projected biopsy darts^63^, concurrent with the collection of photographic, photogrammetry and behavioural data, allowing sampled individuals to be binned to ecotype/morphotype *a priori* to genetic analyses. Most samples were selected from separate collection dates and identified groups (when known) to minimize chances of collecting close relatives or replicate individuals. The sample set included 10 ‘*resident*’, 10 ‘*transient*’, 7 *type B1*, 11 *type B2* and 10 *type C* killer whales.

### DNA extraction, library building and sequencing

DNA was extracted using a variety of common extraction methods, including silica-based filter membranes (Qiagen), standard phenol/chloroform extraction^64^, and lithium chloride^65^. Genomic DNA was then sheared to an average size of ~150-200 bp using a Diagenode Bioruptor NGS. Illumina sequencing libraries were built on the sheared DNA extracts using NEBNext (Ipswich, MA, USA) DNA Sample Prep Master Mix Set 1 following Meyer and Kircher^66^. Libraries were subsequently index amplified for 15 cycles using Phusion High-Fidelity Master Mix (Finnzymes) in 50-μL reactions following the manufacturers guidelines. The libraries were then purified using MinElute PCR purification kit (Qiagen, Hilden, Germany). The DNA concentration of the libraries was measured using a 2100 Bioanalyzer (Agilent Technologies, CA, USA), these were then equimolarly pooled by ecotype and each ecotype pool was sequenced across 5 lanes of an Illumina HiSeq 2000 platform using single read (SR) 100-bp chemistry *(i.e.* a total of 25 lanes).

### Read trimming and mapping

A high quality, 2,249-Mb reference killer whale genome assembly (Oorcal.l, GenBank: ANOL00000000.2, contig N50 size of 70.3 kb, scaffold N50 size of 12.7 Mb)^21^ was used as a mapping reference. For the purpose of this study, the genome was masked for repetitive elements using RepeatMasker^67^ and the Cetartiodactyl repeat library from Repbase^68^. Repetitive elements constitute 41.32% of the killer whale reference genome (929,443,262 sites). A further 80,599 sites were identified as mitochondrial DNA transposed to the nuclear genome (numts) and were masked accordingly. The final assembly was then indexed using BWA v. 0.5.9^69^ to serve as the reference for read mapping.

Illumina HiSeq 2000 reads from each individual were processed with AdapterRemoval^70^ to trim residual adapter sequence contamination and to remove adapter dimer sequences as well as low-quality stretches at 3′ ends. Filtered reads >30 bp were then mapped using BWA v. 0.5.9^69^, requiring a mapping quality greater than 30. Clonal reads were collapsed using the rmdup function of the SAMtools (v. 0.1.18) suite^71^. Ambiguously mapped reads were also filtered out using SAMtools. Consensus sequences were then reconstructed in BAM (Binary sequence/Alignment Map) file format. To ensure all repeat regions, which have the potential to bias population genetic inference, were removed, the per-site coverage was calculated across all 48 individuals. Inspection of the data suggested that sites with a total depth (including data from all 48 individuals) of >200× reads were likely to be unmasked repeats. We therefore further masked these regions, which constituted an additional 0.14% of the genome (3,241,923 sites), resulting in a total masking of 932,685,185 sites (41.46% of the genome).

For illustrative purposes (e.g. Manhattan plots) a synteny based chromosomal assembly of the reference genome was produced by aligning the killer whale scaffolds to a chromosomal assembly of the cow *Bos taurus* (Btau_4.6.1) genome using the *Satsuma* aligner^72^ with default settings. Synteny was conserved and showed no large-scale inter-nor intra-chromosomal rearrangements in any scaffolds (Supplementary Fig. 1). Inferences based on outlier peaks were not influenced by this super-scaffolding process.

Filtered reads were further mapped to a reference mitochondrial genome (GU187176.1) and compared with previously published mitogenome sequences from these individuals^16,73^ The assembled mitogenome sequences were a 100% match with those previously generated for these individuals using targeted sequencing approaches^17,73^. As previously reported^17,73^, the mitogenomes of each ecotype clustered in strongly supported mitochondrial DNA clades, with the exception that one *type B1* individual sampled at a different geographic location to the other *type B1* individuals, had a highly divergent mitogenome haplotype^16^. The relationships among these mitochondrial lineages can be seen in a cladogram of a maximum-likelihood phylogenetic reconstruction (Supplementary Fig. 3a) generated as per reference 74.

### Ancestral state reconstruction

The ancestral state for each site was inferred by mapping whole-genomic sequencing reads of the bottlenose dolphin *(Tursiops truncatus*, Short Read Archive accession code SRX200685)^21,75^ against the killer whale reference genome using BWA read mapper as above. The consensus sequence was called using SAMtools, and ambiguous bases were masked with N’s. Ancestral state could be inferred for 2,206,055,540 (98.1%) of 2,249,565,739 bases.

### Inferring time to most recent common ancestor

Nine of the highest coverage killer whales were selected from our dataset, which included two individuals each of the *resident, transient, type B2* and *type C* ecotypes, and a *type B1* individual. Estimates of the time to most recent common ancestor (TMRCA) were based on the number of derived transitions and number of derived transversions at third codon positions. Exclusively considering mutations at third codon sites was expected to minimize the impact of incomplete purifying selection, which can lead to overestimation of the substitution rate on short timescales^76^. However, some mutations at third codons are non-synonymous, notably more so for transversions than transitions, and putatively ephemeral transversions may therefore result in the overestimation of TMRCA. The lower rate of transversions, compared with transitions is expected to minimize the impact of recurrent mutations at the same site, which could result in an underestimation of TMRCA based on transitions^32^. Therefore, the expectation is that our estimate of TMRCA based on transversions may be upwardly biased and our estimate based on transitions may be downwardly biased.

From a total of 4,781,830 third codon positions (reduced to 3,127,876 when sites with missing data in any of the 9 individuals were masked), 7,547 were inferred to be transversions and 12,784 were inferred to be transitions (with a MAF cut-off of 0.1 so as to exclude potential sequencing errors) from the ancestral state (inferred from comparison with the dolphin genome). Of these, 7,120 transversions and 11,176 transitions were fixed in the killer whale and therefore inferred to have occurred along the branch from the killer whale/bottlenose dolphin ancestor and the MRCA of the killer whales (or be due to incorrect inference of the ancestral state); and 421 transversions and 1,608 transitions occurred within one or more of the killer whale lineages. A further six sites were inferred to have undergone a transversion from the ancestral state in at least one of the killer whale lineages, but had derived transitions in at least one other killer whale lineage. The Ts/Tv ratio of derived mutations at third codon positions was therefore estimated to be 3.8 within the killer whale clade.

The proportion of derived mutations at third codon positions found in one individual and shared in another individual is expected to decrease in comparisons between individuals from populations that diverged longer ago, as the probability that the mutation occurred within just one population following the split increases. The proportion of derived transversions and transitions at third codon positions inferred within the *type B1* individual that were shared with each of the other 8 individuals was measured. The two *resident* individuals shared the least number of derived tranversions with the *type B1* individual (Supplementary Table 3). The results were highly consistent between individuals of the same ecotype (Supplementary Table 3).

The mean rate of nucleotide evolution estimated for odontocetes of 9.10 × 10^−10^ substitutions/site/year (95% HPDI: 6.68 × 10^−10^, 1.18 × 10^−9^)^26^ was then scaled by our estimate of the Ti/Tv ratio of 3.8 at third codon positions within our killer whale dataset and used to predict the time taken to accumulate 124 derived tranversions and 465 transitions at third codon positions that we inferred had been derived in *type B1* since splitting from a shared ancestor with the *resident* ecotype.

### Admixture analysis

An individual-based assignment test was performed to determine whether the ecotypes to which each individual had been assigned *a priori*, based on observed behaviour and/or morphological characteristics at the time of sampling, represented discrete gene pools in Hardy-Weinberg equilibrium. Since the 48 genomes generated for this study had an average sequencing depth of 2×, genotypes can only be called with very high uncertainty. Therefore, NGSadmix^29^, a maximum likelihood method that is based on a model very similar to other maximum likelihood-based admixture methods such as ADMIXTURE^77^ was employed. Whereas all other admixture methods base their inference on called genotypes and implicitly assume that the genotypes are called without error, NGSadmix bases its inference on genotype likelihoods (GLs) and in doing so it takes into account the uncertainty in the called genotypes that is inherently present in low-depth sequencing data^23,24^. The method has been demonstrated, using simulations and publicly available sequencing data, to have great accuracy even for very low-depth data of less than 2-fold mean depth^29^. GLs were estimated using the SAMtools method^71^ implemented in the software package ANGSD^78^. NGSadmix was run with the number of ancestral populations *K* set to 3-5. For each of these *K* values, NGSadmix was re-run multiple times with different seeds in order to ensure convergence. Sites were further filtered to include only autosomal regions covered in at least 40 individuals, and removing sites with a minor allele frequency below 5% estimated from the genotype likelihoods, resulting in the analyses being based upon 603,519 variant sites. The highest likelihood solutions can be seen in Fig. 2C.

### Principal Component Analysis (PCA)

Assignment of individuals to ecotype, and structuring among ecotypes was further investigated using Principal Component Analysis (PCA), implemented in the ngsTools suite^79^ taking genotype uncertainty into account^80^. Briefly, the covariance matrix between individuals, computed as proposed in reference^81^, is approximated by weighting each genotype for its posterior probability, the latter computed using the ANGSD software package^78^ as described in Nielsen *et al*^23^. The eigenvectors from the covariance matrix were generated with the *R* function ‘eigen’ and significance was determined with a Tracy-Widom test to evaluate the statistical significance of each principal component identified by the PCA. The PCA was plotted using an in-house *R* script (available at https://github.com/mfumagalli/ngsPopGen/tree/master/scripts).

### Distance-based phylogenetic inference

We generated 100 matrices of pairwise genetic distances, in which pairwise genetics distances were calculated using ngsDist^82^, which takes genotype uncertainty into account by avoiding genotype calling and instead using genotype posterior probabilities estimated by ANGSD. A block-bootstrapping procedure was used to generate 100 distance matrices, obtained by repetitively sampling blocks of the original data set (Supplementary Table 10). Pairwise genetic distances among the 48 low coverage genomes and high coverage resident genome were visualised as a phylogenetic tree using the distance-based phylogeny inference program FastME 2.0^83^ (Supplementary Fig. 3b).

### Pairwise relatedness estimation

We estimated pairwise relatedness due to identity-by-descent (IBD), i.e. genetic identity due to a recent common ancestor, of every possible combination of two individuals using NgsRelate^84^. NgsRelate provides ML estimates of *R*, where *R* = (*k*_0_,*k*_1_,*k*_2_) and *k*_*m*_ is the fraction of genome in which the two individuals share *m* alleles IBD. NgsRelate provides maximum likelihood estimates of *R* by finding the value of *R* that maximizes this likelihood function with an Expectation Maximization algorithm using genotype likelihoods instead of genotypes to account for the inherent uncertainty of the genotypes. NgsRelate has been shown using simulations and real data to provide robust estimates for low-depth NGS data (as low as 1-2×), which are markedly better than genotype-based methods^84^. The full results are reported in Supplementary Table 11, and a summary is presented in Supplementary Fig. 4.

### TreeMix, *f*3-statistics and D-statistic

The genetic relationships among ecotypes were further reconstructed in the form of a maximum likelihood graph representing the degree of genetic drift generated using TreeMix^31^. TreeMix estimates a bifurcating maximum likelihood tree using population allele frequency data and estimates genetic drift among populations using a Gaussian approximation. The branches of this tree represent the relationship between populations based on the majority of alleles. Migration edges are then fitted between populations that are a poor fit to the tree model, and in which the exchange of alleles is inferred. The addition of migration edges between branches is undertaken in stepwise iterations to maximize the likelihood, until no further increase in statistical significance is achieved^31^. The directionality of gene flow along migration edges is inferred from asymmetries in a covariance matrix of allele frequencies relative to an ancestral population as implied from the maximum likelihood tree. We ran TreeMix fixing the *transient* ecotype as the root, with blocks of 1,000 SNPs (corresponding to approximately several hundred kilobases) to account for linkage disequilibrium among sites. The graphs are presented in Fig. 2a and Supplementary Fig. 5. This method does require genotype calls as an input, and is therefore susceptible to errors associated with genotype calling from low-coverage sequencing data. However, since TreeMix works on population allele frequencies, not genotypes, it was possible to determine the frequencies of the most common alleles with high confidence. The topology is comparable to output from other approaches applied here that do account for genotype uncertainty, providing confidence in the result.

The three-population (*f*3) test, which can provide evidence of admixture, even if gene flow events occurred hundreds of generations ago^34^, was implemented in TreeMix to test for ‘treeness’, i.e. how well relationships can be represented by bifurcations. These tests are of the form *f*3(A;B,C), where a significantly negative value of the *f*3 statistic implies that population A is admixed^34^. *f*3-statistics were computed using the estimators described in reference 34, obtaining standard errors using a block jackknife procedure over blocks of 1,000 SNPs. The results are presented in Supplementary Fig. 6 and Supplementary Tables 4 and 5.

To further investigate the evolutionary history of the killer whale ecotypes, D-statistic (ABBA-BABA) tests^32,33^ were conducted on proposed tree-like histories comprised of all possible combinations of 16 killer whales, with the bottlenose dolphin fixed as the ancestral outgroup. This statistic identifies an excess of shared derived alleles between taxa, which could result from introgression or ancestral population structure. The statistic can thus be used to identify departures from ‘treeness’ of a given topology. For example, if H1, H2 and H3 are taken to denote 3 ecotypes, the test can be used to evaluate if the data are inconsistent with the null hypothesis that the tree (((H1, H2), H3), dolphin) is correct and that there has been no gene flow between H3 and either H1 or H2 or any populations related to them. The definition used here is from reference 33:
D = (*n*ABBA-*n*BABA) / (*n*ABBA+*n*BABA)
where *n*ABBA is the number of sites where H1 shares the ancestral allele with the dolphin, and H2 and H3 share a derived allele (ABBA sites); and, *n*BABA is the number of sites where H2 shares the ancestral allele with the dolphin, and H1 and H3 share a derived allele (BABA sites). Under the null hypothesis that the given topology is the true topology, we expect an equal proportion of ABBA and BABA sites and thus D = 0. Hence a test statistic that differs significantly from 0 provides evidence either of gene flow or the tree being incorrect due to ancestral population structuring. The significance of the deviation from 0 was assessed using a Z-score based on jackknife estimates of the standard deviation of the D-statistics. This Z-score is based on the assumption that the D-statistic (under the null hypothesis) is normally distributed with mean 0 and a standard deviation equal to a standard deviation estimate achieved using the “delete-m Jackknife for unequal m” procedure. The tests were implemented in ANGSD^78^ and performed by sampling a single base at each position of the genome to remove bias caused by differences in sequencing depth.

### Genetic differentiation (*F*_ST_), divergence (*D*_xy_) and nucleotide diversity (*π*)

Measures of genetic differentiation, divergence and diversity were estimated using methods specifically designed for low-coverage sequencing data. Using a Maximum Likelihood-based approach previously proposed^23^ and using the bottlenose dolphin genome to determine the ancestral state of each site, the unfolded site frequency spectrum (SFS) was estimated jointly for all individuals within a population for sites sequenced in ≥5 individuals in each population. Using this ML estimate of the SFS as a prior in an Empirical Bayes approach, the posterior probability of all possible allele frequencies at each site was then computed^23^. For these quantities, the expectations of the number of variable sites and fixed differences between lineages were estimated as the sum across sites of the probability of each site to be variable as previously proposed^85^. Finally, the posterior expectation of the sample allele frequencies was calculated as the basis for further analysis of genetic variation within and between lineages.

*F*_ST_ was estimated with a method-of-moments estimator^86^ based upon both the maximum likelihood estimate of the 2D-SFS^81^ and the sample allele frequency posterior probabilities of the 2D-SFS^81^. The two estimates were highly correlated (Pearson’s correlation coefficient: *r*^2^ > 0.96) for all pairwise comparisons. However, from inspection of the data, the *F*_ST_ estimates generated from sample allele frequency posterior probabilities provided a more accurate estimation of fixed-differences between populations. The likelihood-based method tends to flatten the FST peaks compared to the posterior probabilities method. This can result in masking of *F*_ST_ peaks with increasing genome-wide FST when using the likelihood-based method. Therefore the posterior likelihood estimates are presented in the Manhattan plots of 50-kb sliding windows (Fig. 4a), with further filtering to only include windows for which >10-kb was covered by at least 5 individuals per population (*F*_ST_). We estimated the probability of a site being variable (Pvar). We tried different Pvar cutoffs and counted the number of variable sites at each Pvar. We then decided on Pvar=1 as the number of variable sites matched our expectations from estimates of diversity (*π*) from the two high coverage genomes. We further checked by comparing FST estimates from a recently published whole genome dataset of carrion crows *(Corvis corone)* and hooded crows *(Corvis cornixf*^87^ down-sampled to low coverage. By only considering sites with a Pvar of 1, we obtained FST estimates comparable to the values obtained from the 7-28 × sequences using GATK. Population-specific allele frequencies were estimated with the ancestral state fixed and the derived allele weighted by the probabilities of the 3 possible states. The allele frequencies were estimated based on genotype likelihoods. To assess the robustness of our per-site *F*_ST_ values estimates, we evaluated how well the values correlated with those estimated from RAD-seq data generated in a previous study^25^. We accessed the SNP data in VCF file format generated by the RAD-seq study from Dryad (doi:10.5061/dryad.qk22t) and used VCFtools to calculate per-site *F*_ST_, using sites called at >20× coverage, between 43 individuals of the resident ecotype and 37 individuals of the transient ecotype (Supplementary Fig. 2a). We then performed 1,000 replicates, randomly sub-sampling with replacement ten individuals from each ecotype. The correlation between FST estimates from these random sub-samples ranged from 0.6861 to 0.9372. We found the correlations between estimates of *F*_ST_ from the RAD-seq data and those from our WGS data ranged from 0.5475 to 0.7140 (Supplementary Fig. 2b). The significant correlation in estimates of *F*_ST_ between two different methods using different individuals suggests that these estimates are reliable.

The average number of nucleotide substitutions *D*xy^88^ was then calculated as the mean of pl*q2+p2*ql, where p1 is the allele frequency of allele 1 in population 1 and p2 in population 2, q1 is the allele frequency of allele 2 in population 1 and q2 in population 2. Sites that were not variable in any of the populations were assigned allele frequencies of zero.

Applying the probabilistic model implemented in ANGSD^78^ we estimated the unfolded SFS in steps of 50, 100 and 200-kb windows using default parameters and genotype likelihoods based on the SAMtools error model^71^. From the SFS we derived nucleotide diversity π (Supplementary Table 12). Estimates of nucleotide diversity can be influenced by differences in sequencing coverage and sequencing error. However, it has been shown that using an empirical Bayes approach, implemented in ANGSD^78^, the uncertainties in low-depth data can be overcome to obtain estimates that are similar to those obtained from datasets in which the genotypes are known with certainty^89^. Multiple checks were performed to ensure that estimates of π were not an artefact of the data filtering. Comparable estimates of π were obtained using the method implemented in ANGSD for a single 20× coverage ‘resident’ genome (π = 0.0009) when it was randomly down-sampled to 2× coverage (π = 0.0008). Nucleotide diversity was estimated for sites covered by at least 5 individuals in each population in windows of size 50, 100 and 200-kb (Supplementary Table 12).

### Demographic reconstruction

Pairwise sequentially Markovian coalescent modelling (PSMC) analysis^27^ was performed on the high-coverage diploid autosomal genome sequences of two individuals to investigate changes in effective population (*N*_e_) size. The PSMC model estimates the time to most recent common ancestor (TMRCA) of segmental blocks of the genome and uses information from the rates of the coalescent events to infer *N*_e_ at a given time, thereby providing a direct estimate of the past demographic changes of a population. The method has been validated by its successful reconstructions of demographic histories using simulated data and genome sequences from modern human populations^27^.

A recent study used PSMC to reconstruct ancestral changes in *N*e through time using ~13× and ~20× coverage diploid autosomal genome sequences of a North Atlantic killer whale (accessed ahead of publication by the consortium that generated the data) and a North Pacific killer whale respectively^22^. The authors interpreted the plots as being indicative of a global decline driven primarily by climate change during the last glacial period of the Pleistocene^22^. This study dismissed changes in connectivity having a role in the observed demographic changes as being ‘unlikely to generate the specific pattern observed (strong population decline) or the very similar profiles for each ocean’^22^. However, PSMC heavily relies on the distribution of polymorphic sites across the genome, and in particular, the length of shared runs of homozygosity, and can be biased when heterozygous sites are wrongly called as being homozygous. PSMC plots of genomes with <20× coverage have been shown not to be directly comparable to higher coverage genomes without first applying a correction for an appropriate false negative rate of detecting heterozygotes^90^. Additionally, the effect of mapping short read data to a reference genome comprised of short contigs may also be problematic, especially when many contigs fall below ˜50-kb (the typical size of shared fragment size from 1,000 generations ago in humans), although to the best of our knowledge, the effect of mapping to different quality reference assemblies on PSMC analysis has not been tested to date. Moura *et al.*^22^ mapped short read data from two individual killer whales to a draft assembly of the bottlenose dolphin (assembly turTru1, Ensembl database release 69.1) made up of 0.24 million scaffolds, with a scaffold N50 of 116,287, in which 94% of the scaffolds, comprising approximately 25% of the genome, are less than 50-kb long. Moura *et al.*^22^ thereby generated a 20 × average coverage sequence and a 13 × average coverage sequence, and did not apply a correction for differences in false negative rate of detection of heterozygotes due to the difference in coverage. The PSMC plots from the two genomes presented in Moura *et al.*^22^ do not converge in effective population size (Supplementary Fig. 9) even though the two individuals shared a relatively recent common ancestor (TMRCA estimated at ˜150 KYA by the same authors^28^). In order to better understand if these methodological issues (<20× coverage sequence data mapped to a highly fragmented reference) led to erroneous inference of the demographic histories and the underlying processes in this previous study^22^, PSMC was used to analyse down-sampled versions of the high coverage North Atlantic killer whale genome.

A 50× bam file was produced by mapping the short read data generated from the North Atlantic killer whale to scaffolds of the autosomal regions of the high quality killer whale reference genome that were greater than 10-Mb in length, totaling 1.5 Gb and which had all repeat regions masked as noted above. The 50× coverage bam file was then down-sampled to produce 13× and 20× coverage bam files. An additional 20× bam file was produced for a North Pacific killer whale using data from the short read archive (SRP035610)^22^ and mapped as above. A consensus sequence of each of the four bam files was then generated in fastq format sequentially using: firstly, SAMtools mpileup command with the −C50 option to reduce the effect of reads with excessive mismatches; secondly, bcftools view −c to call variants; lastly, vcfutils.pl vcf2fq to convert the vcf file of called variants to fastq format with further filtering to remove sites with less than a third or more than double the average depth of coverage and Phred quality scores less than 30. The PSMC inference was then carried out using the recommended input parameters^27^ for human autosomal data, i.e. 25 iterations, with maximum TMRCA (*T*_max_) =15, number of atomic time intervals (*n*) = 64 (following the pattern (1*4 + 25*2 + 1*4 + 1*6), and initial theta ratio (*r*) =5. For the initial comparison between the 13 ×, 20× and 50× coverage North Atlantic genomes, a generation time of 25.7 years and a mutation rate of 1.53×10^−8^ substitutions/nucleotide/generation were applied, as per reference 22.

Comparison of the PSMC inference plots based on the 13 ×, 20× and 50× coverage files, generated from the same individual, highlighted the impact of coverage on inference of both the magnitude of *N*_e_ at any given time and the timing of the changes in *N*_e_, consistent with findings by a previous study^90^. In particular, estimates of *N*_e_ in more recent times based on the 13 × genome assembly differed markedly to inferred *N*_e_ from the 20× and 50× genome assemblies (Supplementary Fig. 10). This is a consequence of a higher false negative detection rate of heterozygote sites in the 13 × genome assembly, producing the same effect as a smaller mutation rate would have on the plot. The PSMC plots of the 20× and 50× coverage North Atlantic genome were almost identical both regarding the timing and the magnitude of demographic events. Applying an additional correction for a uniform false detection rate of 2% further increased the similarity of the two plots and suggested that by applying this correction the 20× North Pacific killer whale genome could be directly compared with the 50× North Atlantic killer whale genome using PSMC.

All three plots of the North Atlantic killer whale genome (13×, 20× & 50×) are consistent in estimating a marked decline in *N*_e_ between 100,000 years and 20,000 years ago. To better infer the process underlying this decline in *N*_e_, PSMC was used to compare equal coverage (20×) assemblies of the North Pacific and North Atlantic genomes. As with other inference methods based on coalescent theory, PSMC can only infer scaled times and population sizes. To convert these estimates into real time and size, all scaled results need to be divided by the mutation rate. To allow comparison of the relative timing of population splits with a published time-calibrated nuclear phylogeny based on RAD-seq data we scaled the PSMC plot using the same mutation rate as reference 28. Although the two papers by Moura *et al.*^22,28^, were published almost concurrently, they use two different mutation rates for nuclear genomic data for each analysis: 1.53×10^−8^ substitutions/nucleotide/generation for PSMC^22^ and an estimate almost double this rate for their time-calibrated phylogeny of 2.83×10^−8^ substitutions/nucleotide/generation, based on their given rate of 0.0011 substitutions per site per million years^28^ and a generation time of 25.7 years as above. We therefore scaled the PSMC plots by a generation time of 25.7 years and a mutation rate of 2.83×10^−8^ substitutions/nucleotide/generation. A total number of 100 bootstraps were performed. The combined PSMC plot of both genomes is shown in Supplementary Fig. 11 and compared with population split times from a previously published time-calibrated phylogeny that supports the changes in inferred *N*_e_ by PSMC are at least partially driven by changes in connectivity. This plot was then re-scaled to the mutation rate that we have used for our TMRCA estimation, 2.34×10^−8^ substitutions/nucleotide/generation^26^ and compared to TMRCA of the Atlantic and North Pacific *resident* lineages (estimated as above) and presented in Fig. 3a. Lastly, we simulated data consistent with a population split using ms^91^ and plotted the changes in effective population size inferred by PSMC (Supplementary Fig. 12).

A final PSMC plot of the autosomal regions of the North Atlantic female killer whale at a coverage of 50× was scaled to the autosomal mutation rate (*μ*_A_ of 2.34×10^−8^ substitutions/nucleotide/generation^26^, as used in our estimation of TMRCA (see above) and compared with a plot of the diploid X-chromosome, scaled to real time as per reference 27 in which the neutral mutation rate of the X-chromosome was derived as *μ*X=*μ*A[2(2+α)]/[3(1+α)], assuming a ratio of male-to-female mutation rate of α = 2^92^. This gave us an estimated *μ*X = 2.08×10^−8^ substitutions/nucleotide/generation. The plot is presented in Fig. 3b. We also re-estimated the PSMC plot for the X-chromosome using different mutation rates to investigate which rate would produce PSMC plots with inferred concurrent declines in *N*_e_ in autosomes and X-chromosome. We found that *μ*X = 1.00×10^−8^ substitutions/nucleotide/generation would be needed to synchronise the inferred demographic changes in these two markers (Supplementary Fig. 17). This would require the male-to-female mutation rate (α) to be orders of magnitude higher, making it seemingly biologically unrealistic.

To reconstruct ancestral demography in the ecotypes for which we did not have a high coverage genome, we applied a method that uses the site frequency spectrum from population genomic data to infer ancestral population size changes^44^. The Stairway Plot method first uses site frequency spectrum (SFS) from population genomic sequence data to estimate a series of population mutation rates (i.e. *θ* = 4*N*_e_*μ*, where *μ* is the mutation rate per generation and *N*_e_ is the effective population size), assuming a flexible multi-epoch demographic model. Changes in effective population size through time are then estimated based on the estimations of *θ* As input data, we transformed the probability estimates of our site frequency spectra into SNP counts. We first ran the method on the resident ecotype to compare with the demographic reconstruction suggested by the PSMC analysis on the high covered North Pacific individual (see above). Population structure is a notable confounding factor for inferring demographic history^41,93,94^. Consistent with this, we estimate broad confidence intervals of our estimates of *N*_e_ subsequent to the estimate bottlenecks in each ecotype, during a period overlapping with previous estimates of within-ecotype population splits^28^. As we have sampled individuals from multiple sub-populations of the *resident* and *transient* (and possibly the Antarctic types, although less in known about population structuring in Antarctic waters), we potentially skew the SFS toward low-frequency polymorphism, thereby mimicking the pattern generated by population expansion^93,94^. Previous studies have estimated that subpopulations of the *resident* ecotype split during the last ~10,000 years based in IMa analysis of microsatellite data^38^, and during the last ~80,000 years based on BEAST analysis of RAD-seq data^28^. Similarly, these same studies estimated that subpopulations of the *transient* ecotype split during the last ~12,000 years based in IMa analysis of microsatellite data^38^, and during the last ~120,000 years based on BEAST analysis of RAD-seq data^28^. We therefore opted not to include SNP counts for singletons and doubletons, which are expected to have arisen recently within each ecotype and not had time to be shared throughout the population, as these may be biased by our sampling protocol and low coverage sequencing data, and because our interest was in demographic change during population splits which were estimated to have occurred over 10,000 years ago. Our focus is on the timing and extent of the bottlenecks within each ecotype, which all individuals within an ecotype coalesce back to and therefore predate within-ecotype population splits and sub-structuring and the emergence of derived singleton and doubletons.

The inference of demographic history from population genomic data by the Stairway Plot method provided a somewhat comparable result to reconstruction from a single 20× genome by PSMC in respect to the time of onset and magnitude of a demographic decline inferred by both methods. The Stairway Plot method inferred a sudden drop in *N*e, whereas a more gradual decline was inferred by PSMC, consistent with simulations showing that PSMC can infer abrupt changes in *Ne* as gradual changes^27^. The sudden change in estimated effective population size in the stairway plot is due to the method being based on a multi-epoch method, in which epochs coincide with coalescent events^44^. Therefore, the plot is not continuous, but rather it depicts discrete blocks of time (epochs). The number of epochs is determined by the number of individuals within each sample and the number of SNP bins used *i.e.* the number of possible coalescent events. The Stairway Plot method inferred a subsequent and rapid expansion, whereas PSMC did not infer an expansion within our cut-off point of 10,000 years ago, but did infer a more gradual expansion during the Holocene. It should be kept in mind that the PSMC plot is based on data from a single individual and so will track declines in *Ne* due to further founder effects as the resident ecotype continues to split into multiple discrete populations. In contrast, the Stairway Plot is based on population genomic data and will track the change in Ne across all the sampled resident populations after they have split.

The method was then applied to the site frequency spectra of the transient ecotype and type C. The results are shown in Fig. 3c, and in each case the Stairway Plot infers a sudden and dramatic demographic decline, consistent with previously inferred population split times followed by a demographic expansion. The timing of the decline of the Antarctic types overlapped due to the recent shared ancestry and so only the plot for type C is shown in Fig. 3c for clarity.

### Inferring putatively functional allele shifts due to selection

Shifts in allele frequencies can occur due to selection, but differences in allele frequencies can also accumulate between populations due to drift^45,46^. To infer shifts in allele frequencies potentially due to selection we considered the 1% of SNPs with the highest *F*_ST_ values from each pairwise comparison between killer whale ecotypes. However, as *F*_ST_ is dependent on the underlying diversity of the locus even extreme outlier loci can be due to genetic drift alone. We therefore additionally looked for over-representation of the top 1% outliers in different categories, e.g. exons, 25-kb flanking regions (potential regulatory regions), introns and intergenic regions, at different *F*_ST_ bins using a chi-square test. Residuals are expected to be normally distributed and indicate statistical significance of over-and under-representation of specific categories. The significance threshold was subjected to Bonferroni correction. The top 100 outliers in exons from each pairwise comparison were then used for gene ontology (GO) overrepresentation analysis^95^ to identify enrichment due to diet (in mammal-versus fish-eating ecotypes) and climate (in Antarctic versus Pacific).

To more robustly infer whether genetic changes in exons were associated with ecotype divergence due to selection we applied the population branch statistic (PBS)^51^. The PBS has strong power to detect (even incomplete) selective sweeps over short divergence times^5,51,96^, making it relevant for the scenario we are investigating in this study. We therefore estimated the PBS for 50-kb sliding windows shifting in 10-kb increments (approximating to a window size of 10 SNPs) to detect regions of high differentiation potentially due to selective sweeps. We followed the approach of Yi *et al.*^51^, and used the classical transformation by Cavalli-Sforza^97^,

*T*=-log(l -*F*_ST_) to obtain estimates of the population divergence time *T* in units scaled by the population size. For each 50-kb window, we calculated this value between each pair of ecotypes. The length of the branch leading to the one ecotype since the divergence from a recent ancestor (for example in the equation given the length of the branch to type B1 since diverging from types B2 and C) is then obtained as

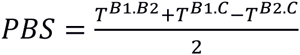

These window-based PBS values represent the amount of allele frequency change at a given 50-kb genomic region in the history of this ecotype (since its divergence from the other two populations)^51^. To further narrow down the target of selection, the PBS was estimated for exons as per reference 5 and compared to genome-wide values to identify if they were in the top 99.9 percentile (for the branches to the *resident, transient* and ancestral *Antarctic* ecotypes) and the top 99.99 percentile for the shorter branches leading to each Antarctic ecotype from their most recent common shared ancestor. We further filtered outliers to just include exons that encoded non-synonymous amino acid substitutions and searched the database GeneCards^98^ for functions of the encoded proteins to identify potential targets of natural selection due to ecological differences. Supplementary Table 13 contains a list of outlier loci and the corresponding PBS value for each branch. Supplementary Table 14 contains details of fixed non-synonymous changes in the exons of protein coding genes, including the per-individual and per-population read count for each allele.

## Accession codes

All sequencing data are available in European Nucleotide Archive (ENA) under accession numbers ERS554424- ERS554471.

Note: Any Supplementary Information and Source Data files are available in the online version of the paper.

## ACKNOWLEDGEMENTS

The project was funded by a Marie Curie IEF ‘KWAF10‘ and Lawksi Foundation grants to ADF; European Research Council grant ERCStG-336536 to JBWW; a Danish National Research Foundation grant DNRF94 to MTPG; by Swiss SNSF Grant 31003A-143393 to LE and by a short visit grant to ADF from the ESF Research Networking Programme ConGenOmics. We would like to acknowledge The Danish National High-Throughput DNA Sequencing Centre for sequencing the samples and in particular to thank Andaine Seguin-Orlando, Lillian Petersen, Cecilie Demring Mortensen, Kim Magnussen and Ian Lissimore for technical support. Large-scale computational effort was made possible by the UPPMAX next-generation sequencing cluster and storage facility (UPPNEX), funded by the Knut and Alice Wallenberg Foundation and the Swedish National Infrastructure for Computing. Heng Li, Richard Durbin, Line Skotte, Anders Albrechtsen, Stephan Schiffels, Lounès Chikhi and Rasmus Nielsen provided useful feedback and discussions on the methods. We thank Willy Rodríguez for providing scripts for the ms simulations. Mike Ford and Kim Parsons provided useful feedback on an earlier draft of this manuscript. We thank Paul Fiedler for providing the map used in figure 1 and Uko Gorter for providing the illustrations of killer whale ecotypes and Robert Pitman’s support and input throughout this project. We are grateful to M. Ford, K. Parsons, A. Burdin, M. Dahlheim, R. Pitman, the SWFSC Marine Mammal and Sea Turtle Research Collection, and to the International Whaling Commission for providing samples.

*We dedicate this work to the memory of Eva Saulitis, whose rich legacy includes an invaluable contribution to the foundations of our understanding of killer whale biology, without which this study would not have been possible*.

## AUTHOR CONTRIBUTIONS

A.D.F., P.A.M. and M.T.P.G. initially conceived and designed the study, which was further developed by J.B.W.W., N.V. and input from all other authors; R.W.B., J.W.D., B.H. and P.W. conducted field work and collected biopsy samples; K.M.R. and P.A.M. conducted DNA extraction and sample selection; A.D.F. conducted DNA library construction; A.D.F. and M.C.A.-A. conducted sequence read filtering and mapping; N.V., A.D.F., F.G.V. M.F., M.D.M., T.S.K., T.V. and V.C.S. ran the population genetic analyses; A.D.F. and N.V. wrote the manuscript and the Supplementary Information with input from J.B.W.W., M.T.P.G. and the other authors.

## References

1. Laland, K. N., Odling-Smee, J. & Myles, S. How culture shaped the human genome: bringing genetics and the human sciences together. Nature Rev. Genet. 11, 137–148 (2010).

2. Wang, E. T., Kodama, G., Baldi, P. & Moyzis, R. K. Global landscape of recent inferred Darwinian selection for Homo sapiens. Proc. Natl Acad. Sci. USA 103, 135–140 (2006).

3. Hawks, J. Wang, E. T., Cochran, G. M., Harpending, H. C. & Moyzis, R. K. Recent acceleration of human adaptive evolution. Proc. Natl. Acad. Sci. U.S.A. 104, 20753–20758 (2007).

4. Moltke, I. et al. A common Greenlandic TBC1D4 variant confers muscle insulin resistance and type 2 diabetes. Nature 512, 190–193 (2014).

5. Fumagalli, M. et al. Greenlandic Inuit show genetic signatures of diet and climate adaptation. Science 349, 1343–1347 (2015).

6. Varki, A., Geschwind, D. H. & Eichler, E. E. Explaining human uniqueness: genome interactions with environment, behaviour and culture. Nature Rev. Genet. 9, 749–763 (2008).

7. Ford, J. K. B. in The Encyclopedia of Marine Mammals, 2nd edn, (eds Perrin, W. F., Würsig, B. & Thewissen, J. G M.) 650–657 (Elsevier, 2009).

8. Ford, J. K. B. et al. Dietary specialization in two sympatric populations of killer whale (Orcinus orcd) in coastal British Columbia and adjacent waters. Can. J. Zool. 76, 1456–1471 (1998).

9. Saulitis E. L. et al. Foraging strategies of sympatric killer whale (Orcinus orcd) populations in Prince William Sound, Alaska. Mar. Mamm. Sci. 16, 94–109 (2000).

10. Matkin C. O. et al. Ecotypic variation and predatory behavior among killer whales (Orcinus orcd) off the eastern Aleutian Islands, Alaska. Fish. Bull. 105, 74–87 (2007).

11. Filatova O. A. et al. Reproductively isolated ecotypes of killer whales Orcinus orca in the seas of the Russian Far East. Biology Bulletin 42, 674–681 (2015).

12. Pitman, R. L. & Ensor, P. Three forms of killer whales (Orcinus orcd) in Antarctic waters. J. Cetacean Res. Manage. 5, 131–139 (2003).

13. Pitman R. L. & Durban, J. W. Killer whale predation on penguins in Antarctica. Polar Biol. 33, 1589–1594(2010).

14. Pitman R. L. & Durban, J. W. Cooperative hunting behavior, prey selectivity and prey handling by pack ice killer whales (Orcinus orca), type B, in Antarctic Peninsula waters. Mar. Mamm. Sci. 28, 16–36 (2012).

15. Durban, J. W., Fearnbach, H, Burrows, D. G., Ylitalo, G.M. & Pitman, R.L. Morphological and ecological separation of Type B killer whales around the Antarctic Peninsula. Polar Biol (in review).

16. Morin, P. A. et al. Geographic and temporal dynamics of a global radiation and diversification in the killer whale. Mol. Ecol. 24, 3964–3979 (2015).

17. Brent, L. J. N. et al. Ecological knowledge, leadership, and the evolution of menopause in killer whales. Curr. Biol. 25, 1–5 (2015).

18. Riesch, R., Barrett-Lennard, L. G, Ellis, G M., Ford, J. K. B. & Deecke, V. B. Cultural traditions and the evolution of reproductive isolation: ecological speciation in killer whales? Biol. J. Linn. Soc. 106, 1–17 (2012).

19. Whitehead, H. Cultural selection and genetic diversity in matrilineal whales. Science 282, 1708–1711 (1998).

20. Laland, K. N. & Janik, V. M. The animal cultures debate. Trends Ecol. Evol. 21, 542–547 (2006).

21. Foote, A. D. et al. Convergent evolution of marine mammal genomes. Nature Genetics, 47, 212–215 (2015).

22. Moura A. E. et al. Killer whale nuclear genome and mtDNA reveal widespread population bottleneck during the Last Glacial Maximum. Mol. Biol. Evol. 31, 1121–1131 (2014).

23. Nielsen R., Korneliussen, T., Albrechtsen, A., Li, Y. & Wang, J. SNP calling, genotype calling, and sample allele frequency estimation from new-generation sequencing data. PloS one 7, e37558 (2012).

24. O’Rawe, J. A., Ferson, S. & Lyon, G. J. Accounting for uncertainty in DNA sequencing data. Trends Genet. 31, 61–66 (2015).

25. Moura, A. E. et al. Population genomics of the killer whale indicates ecotype evolution in sympatry involving both selection and drift. Mol. Ecol. 23, 5179–5192(2014).

26. Dornburg A., Brandley, M. C, McGowan, M. R. & Near, T. J. Relaxed clocks and inferences of heterogeneous patterns of nucleotide substitution and divergence time estimates across whales and dolphins (Mammalia: Cetacea). Mol. Biol. Evol. 29, 721–739(2011).

27. Li, H. & Durbin, R. Inference of human population history from individual whole-genome sequences. Nature 475, 493–496 (2011).

28. Moura, A. E. et al. Phylogenomics of the killer whale indicates ecotype divergence in sympatry. Heredity, 114, 48–55 (2015).

29. Skotte, L., Sand Korneliussen, T. & Albrechtsen, A. Estimating individual admixture proportions from Next Generation Sequencing data. Genetics 195, 693–702 (2013).

30. Seehausen, O. et al. Genomics and the origin of species. Nature Rev. Genet. 15, 176–192 (2014).

31. Pickrell, J. K. & Pritchard, J. K. Inference of population splits and mixtures from genome-wide allele frequency data. PLoS Genet. 8, e1002967 (2012).

32. Green R. E. et al. A draft sequence of the Neandertal genome. Science 328, 710–722 (2010).

33. Durand, E. Y., Patterson, N, Reich, D. & Slatkin, M. Testing for ancient admixture between closely related populations. Mol. Biol. Evol. 28, 2239–2252 (2011).

34. Patterson, N et al. Ancient admixture in human history. Genetics 192, 1065–1093 (2012).

35. Sousa, V., & Hey, J. Understanding the origin of species with genome-scale data: modelling gene flow. Nature Rev. Genet. 14, 404–414 (2013).

36. Nei, M., Maruyama, T. & Chakraborty, R. The bottleneck effect and genetic variability in populations. Evolution 29, 1–10 (1975).

37. Marth, G. T., Czabarka, E., Murvai, J. & Sherry, S. T. The allele frequency spectrum in genome-wide human variation data reveals signals of differential demographic history in three large world populations. Genetics 166, 351–372 (2004).

38. Hoelzel, A. R. Evolution of population structure in a highly social top predator, the killer whale. Mol. Biol. Evol. 76, 1407–1415 (2007).

39. Pilot, M., Dahlheim, M. E. & Hoelzel, A. R. Social cohesion among kin, gene flow without dispersal and the evolution of population genetic structure in the killer whale (Orcinus orca). J. Evol. Biol. 23, 20–31 (2010).

40. Lanfear, R., Kokko, H. & Eyre-Walker, A. Population size and the rate of evolution. Trends Ecol. Evol. 29, 33–41 (2014).

41. Mazet, O., Rodríguez W., Grusea, S., Boitard, S. & Lounès, C. On the importance of being structured: instantaneous coalescence rates and human evolution-lessons for ancestral population size inference? Heredity doi:10.1038/hdy.2015.104 (2015).

42. Pool, J. E. & Nielsen, R. Population size changes reshape genomic patterns of diversity. Evolution Int. J. Org. Evolution 61, 3001–3006 (2007).

43. Keinan, A., Mullikin, J. C, Patterson, N. & Reich, D. Accelerated genetic drift on chromosome X during the human dispersal out of Africa. Nature Genet. 41, 66–70 (2009).

44. Liu, X. & Fu, Y.-X. Exploring population size changes using SNP frequency spectra. Nature Genetics, 47, 555–559 (2015).

45. Excoffier, L., Foll, M. Petit, R. J. Genetic consequences of range expansions. Annu. Rev. Ecol. Evol. Syst. 40, 481–501 (2009).

46. Kimura M., Ohta, T. The average number of generations until fixation of a mutant gene in a finite population. Genetics 61, 763–771 (1969).

47. Cruickshank, T. E. & Hahn, M. W. Reanalysis suggests that genomic islands of speciation are due to reduced diversity, not reduced gene flow. Mol. Ecol. 23, 3133–3157(2014).

48. Burri, R. et al. Linked selection and recombination rate variation drive the evolution of the genomic landscape of differentiation across the speciation continuum of Ficedula flycatchers. Genome Res. 25, 1656–1665 (2015).

49. Swanson, W. J. & Vacquier, V. D. Rapid evolution of reproductive proteins. Nat. Rev. Genet. 3, 137–144 (2002).

50. Liu S. et al. Population genomics reveal recent speciation and rapid evolutionary adaptation in polar bears. Cell 157, 785–792 (2014).

51. Yi et al. Sequencing of 50 human exomes reveals adaptation to high altitude. Science 329, 75–78 (2010).

52. Forman, O. P., Pendens, J., Hartley, C, Hayward, L. J., Ricketts, S. L. & Mellersh, C.S. Parallel mapping and simultaneous sequencing reveals deletions in BCAN and FAM83H associated with discrete inherited disorders in a domestic dog breed. PLoS Genet. 8, e1002462 (2012).

53. Springer, M. S., Starrett, J., Morin, P. A., Lanzetti, A., Hayashi, C. & Gatesy, J. Inactivation of C4orf26 in toothless placental mammals. Mol. Phyl. Evol. 95, 34–45 (2016).

54. Kuga, T. et al. A novel mechanism of keratin cytoskeleton organization through casein kinase Iα and FAM83H in colorectal cancer. J. Cell Science 126, 4721–4731 (2013).

55. Loschke, F., Seltmann, K., Bouameur, J. E. & Magin, T. M. Regulation of keratin network organization. Curr. Opin. Cell. Biol. 32, 56–64 (2015).

56. Durban, J. W. & Pitman R. L. Antarctic killer whales make rapid, round-trip movements to subtropical waters: evidence for physiological maintenance migrations? Biol. Lett. 8, 274–277 (2012).

57. Satoh, T. & Hosokawa, M. The mammalian carboxylesterases: from molecules to functions. Ann. Rev. Pharmacol. Toxicol. 38, 257–288 (1998).

58. Finkelstein, J. D. Methionine metabolism in mammals. J. Nutr. Biochem. 1, 228–237 (1990).

59. Mann, N. J, Li, D., Sinclair, A. J., Dudman, N. P. B., Guo, X. W., Elsworth, G R, Wilson, A. K. & Kelly, F. D. The effect of diet on plasma homocysteine concentrations in healthy male subjects. Eur. J. Clin. Nutr. 53, 895–899 (1999).

60. Hamm, D., Mautz, B. S., Wolfner, M. F., Aquadro, C. F. & Swanson, W. J. Evidence of amino acid diversity-enhancing selection within humans and among primates at the candidate sperm-receptor gene PKDREJ. Am. J. Hum. Genet. 81, 44–52 (2007).

61. Powell, A., Shennan, S. & Thomas, M. G. Late Pleistocene demography and the appearance of modern human behavior. Science 324, 1298–1301 (2009).

62. Atkinson, Q. Phonemic diversity supports a serial founder effect model of language expansion from Africa. Science 332, 346–348 (2011).

63. Barrett-Lennard, L. G., Smith, T. G. & Ellis, G. M. A cetacean biopsy system using lightweight pneumatic darts and its effect on the behavior of killer whales. Mar. Mamm. Sci. 12, 14–27 (1996).

64. Sambrook, J. Fritsch, E. F. & Maniatis, T. Molecular cloning: a laboratory manual. (Cold Spring Harbor Laboratory Press, 1989).

65. Gemmell, N. J. & Akiyama, S. An efficient method for the extraction of DNA from vertebrate tissues. Trends Genet. 12, 338–339 (1996).

66. Meyer, M. & Kircher, M. Illumina sequencing library preparation for highly multiplexed target capture and sequencing. Cold Spring Harbor Protocols 6 doi:10.1101/pdb.prot5448 (2010)

67. Smit, A., Hubley, R. & Green, P. RepeatMasker Open-3.0. www.repeatmasker.org (1996).

68. Jurka, J. Kapitonov, V. V., Pavlicek, A., Klonowski, P., Kohany, O. & Walichiewicz, J. Repbase Update, a database of eukaryotic repetitive elements. Cytogenet. Genome Res. 110, 462–467 (2005).

69. Li, H. & Durbin, R. Fast and accurate short read alignment with Burrows-Wheeler transform. Bioinformatics 25, 1754–1760 (2009).

70. Lindgreen, S. AdapterRemoval: easy cleaning of next-generation sequencing reads. BMC Res. Notes, 5, 337 (2012).

71. Li, H. et al. The sequence alignment/map format and SAMtools. Bioinformatics, 25, 2078–2079 (2009).

72. Grabherr, M. G. et al. Genome-wide synteny through highly sensitive sequence alignment: Satsuma. Bioinformatics 26, 1145–1151 (2010).

73. Morin, P. A. et al. Complete mitochondrial genome phylogeographic analysis of killer whales (Orcinus orcd) indicates multiple species. Genome Res. 20, 908–916 (2010).

74. Foote A. D. et al. Tracking niche variation over millennial timescales in sympatric killer whale lineages Proc. R. Soc. B 280, 20131481 (2013).

75. Lindblad-Toh K. et al. A high-resolution map of human evolutionary constraint using 29 mammals. Nature 478, 476–482 (2011).

76. Ho, S. Y. W. et al. Time-dependent rates of molecular evolution. Mol. Ecol. 20, 3087–3101 (2011).

77. Alexander, D. H, Novembre, J. & Lange, K. Fast model-based estimation of ancestry in unrelated individuals. Genome Res. 19, 1655–1664 (2009).

78. Korneliussen, T. S., Albrechtsen, A. & Nielsen, R. ANGSD: Analysis of Next Generation Sequencing Data. BMC Bioinformatics 15: 356 (2014).

79. Fumagalli, M., Vieira F. G., Linderoth, T. & Nielsen, R. ngsTools: methods for population genetics analyses from next-generation sequencing data. Bioinformatics 30, 1486–1487 (2014).

80. Fumagalli M. et al. Quantifying population genetic differentiation from next generation sequencing data. Genetics 195, 979–992 (2013).

81. Patterson, N., Price, A. L. & Reich, D. Population structure and eigenanalysis. PLoS Genetics, 2, e190 (2006).

82. Vieira, F. G., Lassalle, F., Korneliussen, T. S. & Fumagalli, M. Improving the estimation of genetic distances from Next-Generation Sequencing data. Biol. J. Linnean Soc. 117, 139–149 (2015).

83. Lefort, V., Desper, R. & Gascuel, O. FastME 2.0: a comprehensive, accurate, and fast distance-based phylogeny inference program. Mol. Biol. Evol. 32, 2798–2800 (2015).

84. Korneliussen, T. S. & Moltke I. NgsRelate: a software tool for estimating pairwise relatedness from next-generation sequencing data. Bioinformatics 31, 4009–4011.

85. Fumagalli, M. Assessing the effect of sequencing depth and sample size in population genetics inferences. PLoS One 8, e79667 (2013).

86. Reynolds, J., Weir, B. S. & Cockerham, C. C. Estimation of the coancestry coefficient: basis for a short-term genetic distance. Genetics, 105, 767–779 (1983).

87. Poelstra, J. W., Vijay, N., Bossu, C. M., Lantz, H., Ryll, B., Muller, L, Baglione, V., Unneberg, P., Wikelski, M., Grabherr, M. G. & Wolf, J. B. W. The genomic landscape underlying phenotypic integrity in the face of gene flow in crows. Science 344, 1410–1414 (2014).

88. Nei, M. Molecular Evolutionary Genetics (Columbia Univ. Press, 1987).

89. Korneliussen, T. S. et al., Calculation of Tajima’s D and other neutrality test statistics from low depth next-generation sequencing data. BMC Bioinformatics 14, 289 (2013).

90. Orlando, L. et al. Recalibrating Equus evolution using the genome sequence of an early Middle Pleistocene horse. Nature 499, 74–78 (2013).

91. Staab, P. R., Zhu, S., Metzler, D. & Lunter, G. scrm: efficiently simulating long sequences using the approximated coalescent with recombination. Bioinformatics 31, 1680–1682(2015).

92. Miyata, T., Hayashida, H., Kuma, K, Mitsuyasu, K. & Yasunaga, T. Male-driven molecular evolution: a model and nucleotide sequence analysis. Cold Spring Harb. Symp. Quant. Biol. 52, 863–867 (1987).

93. Gattepaille, L. M., Jakobsson, M. & Blum, M. G. B. Inferring population size changes with sequence and SNP data: lessons from human bottlenecks. Heredity 110, 409–419(2013).

94. Ptak, S. & Przeworski, M. Evidence for population growth in humans is confounded by population structure. Trends Genet. 18, 559–563 (2002).

95. Huang, D. W., Sherman, B. T. & Lempicki, R. A. Systematic and integrative analysis of large gene lists using DAVID bioinformatics resources. Nature protocols 4, 44–57 (2008).

96. Zhan, S. et al. The genetics of monarch butterfly migration and warning colouration. Nature 514, 317–321 (2014).

97. Cavalli-Sforza, L. L. Human Diversity. Proc. 12th Int. Congr. Genet. 2, 405–416 (1969).

98. Rebhan M., Chalifa-Caspi, V., Prilusky, J. & Lancet, D. GeneCards: a novel functional genomics compendium with automated data mining and query reformulation support. Bioinformatics 14, 656–664 (1998).

99. Locasale, J. W. Serine, glycine and one-carbon units: cancer metabolism in full circle. Nat. Rev. Cancer 13, 572–583 (2013).

100. Shao, Z. et al. Crystal structure of tRNA m^1^G9 methyltransferase TrmlO: insight into the catalytic mechanism and recognition of tRNA substrate. Nucleic Acids Res. 42, 509–525 (2014).

